# Single-cell transcriptomics reveal alveolar macrophages-specific responses in single-hit ozone exposure model in mice

**DOI:** 10.1101/2025.07.11.664370

**Authors:** Thao Vo, Ishita Choudhary, Sonika Patial, Yogesh Saini

## Abstract

Alveolar macrophages (AMs), a highly plastic immune cell population, are among the first responders to the inhaled ozone (O_3_) and ozonized products in the lung airspaces. However, the complete understanding of how AMs respond to O_3_, particularly to different concentrations, remains elusive. Accordingly, we exposed C57BL/6J male adult mice to filtered air (FA), 1 ppm O_3_, or 1.5 ppm O_3_ for 3 hours. As compared to FA-exposed mice, O_3_-exposed mice exhibited increased recruitment of alveolar macrophages and neutrophils in the lung airspaces, which was consistent with the elevated levels of macrophage- and neutrophil-specific chemokines, i.e., MIP-2, MCP-3, and MCP-5. Next, to profile AM transcriptome from O_3_-exposed mice and understand the relevance of these transcriptomic changes in relation to their population heterogeneity and functionality, we performed single-cell RNA sequencing (scRNA-seq) analyses. The differentially expressed genes (DEGs) analyses on AM population identified significant changes in 1 ppm-exposed and 1.5 ppm O_3_-exposed mice. As compared to AMs from FA-exposed group, AMs from 1 ppm O_3_- and 1.5 ppm O_3_-exposed groups displayed enrichment of pathways including oxidative phosphorylation, EIF2 signaling, and non-canonical NF-kB signaling. Furthermore, AMs from 1 ppm O_3_-exposed mice showed enrichment of IL-10 signaling pathway. On the other hand, AMs from 1.5 ppm O_3_-exposed mice were uniquely enriched in DNA damage bypass and repair pathways. Interestingly, UMAP analyses on annotated AMs resulted in five distinct subclusters. DEGs and ingenuity pathways (IP) analyses on each subcluster revealed O_3_ concentration-dependent enrichment of pathways relevant to protein translation, cholesterol biosynthesis and mitochondrial biogenesis. Further analyses revealed that O_3_ exposure results in cluster-specific alterations to the expression of gene signatures associated with macrophage activation. Finally, AMs from 1.5 ppm O_3_-exposed mice displayed elevated expression of proliferation-associated gene signatures. Taken together, this study identifies O_3_ concentration-dependent alterations in AMs transcriptome and associated functional modulations at a single-cell resolution.

## Introduction

Ground-level ozone (O_3_) is a highly reactive air pollutant generated through photochemical reactions of nitrogen oxides and volatile organic compounds in the presence of heat and ultraviolet sunlight (Aucamp, 2007). O_3_ interacts with the constituents of the epithelial lining fluid (ELF) to generate harmful ozonated products (Frampton et al., 1999b, 1999a; Kelly & Mudway, 2003; Kotiaho et al., 2000; Mudd et al., 1969; Pryor et al., 1996; Pryor & Uppu, 1993). O_3_ causes adverse effects on respiratory health, including altering airway epithelial homeostasis, increasing airway inflammation, and altering lung function (Bell et al., 2004; Jang et al., 2002; Kehrl et al., 1987; C. S. Kim et al., 2011; Mar & Koenig, 2009). Elevated levels of ground-level O_3_ contribute to a significant increase in hospitalization, resulting in substantial health and economic burden (Lin et al., 2008; Medina-Ramón et al., 2006; Moore et al., 2008; Schwartz, 1994).

Alveolar macrophages (AMs) represent a highly plastic and heterogeneous population of sentinel cells that patrol the lung airspaces (Dick et al., 2022; Hussell & Bell, 2014; Wu et al., 2023; Xu-Vanpala et al., 2020). Due to the presence of multiple surface receptors, including Toll-like receptors, scavenger receptors, C-type lectin receptors, and cytokine receptors, AMs are specialized in sensing the presence of both biotic and abiotic entities in the lung airspaces and mount a wide array of functional responses, including antigen presentation, pathogen killing, tissue destruction, tissue repair, and scavenging (Ley et al., 2016; Taylor et al., 2005; Wynn & Vannella, 2016). AMs are likely the first cell type to interact with O_3_ and ozonated products (Patial & Saini, 2020) and subsequently to respond in a variety of ways, including functional activation, release of inflammatory mediators, and communication with neighboring cells (Mosser & Edwards, 2008; Choudhary, Vo, Paudel, Patial, et al., 2021; Choudhary, Vo, Paudel, Wen, et al., 2021). However, the effect of single exposure of O_3_ on resident AMs, at a single-cell resolution, remains unclear.

In this study, we hypothesized that the concentration-dependent effects of O_3_ on AM functional heterogeneity can be identified by evaluating their transcriptomic signatures a single-cell resolution. To test our hypothesis, we performed single-cell RNA sequencing (scRNA-seq) on bronchoalveolar lavage (BAL) cells from C57BL/6J mice that were exposed to either filtered air (FA), 1 ppm O_3_, or 1.5 ppm O_3_. Specifically, we performed gene expression profiling and biological pathways enrichment analyses on AM clusters observed in mice exposed to FA, 1 ppm O_3,_ or 1.5 ppm O_3_. The results from this study highlight the heterogeneity of AMs and the concentration-dependent effects of O_3_ exposure on cluster-specific AMs proliferation, functional activation, and associated enriched biological pathways.

## Materials and Methods

### Animal husbandry

12-week-old male C57BL/6J mice were procured from Jackson Laboratory (Bar Harbor, ME). Mice were allowed to acclimatize for a period of 1 week upon arrival at North Carolina State University Laboratory Animal Resource (NCSU-LAR). Mice were maintained in individually ventilated, hot-washed cages on a 12h:12h dark-light cycle and were fed a regular diet and water *ad libitum*. All animal procedures were approved by the NCSU Institutional Animal Care and Use Committee (IACUC).

### Experimental design and filtered air/ozone exposure

C57BL/6J mice were randomly assigned to their designated exposure groups, i.e., filtered air (FA), 1 ppm ozone (O_3_), or 1.5 ppm O_3_. Mice were transferred into steel wire mesh cages and were placed inside the light-protected chambers without food and water before the start of LAR night cycle. Mice were exposed to FA, 1 ppm O_3_, or 1.5 ppm O_3_ in the designated chambers for 3 hours. O_3_ was generated from the Ozone Generator (TSE, Chesterfield, MO) and the real-time O_3_ concentration in the chamber was monitored using a UV photometric O_3_ analyzer (EnviaAltech Environment, Geneva, IL). Data, including the O_3_ concentration, along with chamber temperature and pressure, were recorded using DACO monitoring and control software (TSE Systems, Chesterfield, MO) throughout the exposure.

### Bronchoalveolar lavage fluid collection

Mice exposed to FA or O_3_ were euthanized 22-24 h after the end of the exposure. Briefly, mice were anesthetized via intraperitoneal injection of 2,2,2-tribromoethanol (Millipore Sigma, Burlington, MA). Midline laparotomy was performed to expose and severe inferior vena cava for exsanguination. Thereafter, thoracotomy was performed to expose the lung and trachea. The left main stem bronchus was ligated, and the right lung lobes were lavaged with a calculated volume of ice-cold Dulbecco’s Phosphate Buffered Saline (DPBS) (Corning, Manassas, VA). The retrieved bronchoalveolar lavage fluid (BALF) was centrifuged at 500 x *g* for 5 min at 4°C. Cell-free supernatant was stored at -80°C for cytokine analyses. After resuspending the cell pellet in 500 μl of DPBS, 90 μl of cell suspension was diluted in 110 μl of DPBS and was used to prepare cytospins. Prepared cytospins were differentially-stained (Wright Giemsa Stain Kit; Newcomer Supply, Middleton, WI) and analyzed to determine the composition of various immune cells. The photographs were captured under the 40X objective of the ECLIPSE Ci-L microscope with DS-Fi2 camera attachment (Nikon, Melville, NY). To maximize the recovery of airspace immune cells, further lavages were performed to collect an additional 5 ml of BALF. The remaining portion of the immune cells from the first lavage and the additional 5 ml lavage of 3-4 mice per group were pooled and centrifuged at 500 x *g* for 5 min at 4°C. The supernatant was removed, and the cell pellet was resuspended in 500 μl of fresh DPBS. The washing process was repeated one more time. After the last wash, the BALF was centrifuged at 500 x *g* for 5 min at 4°C, the supernatant was removed, and the cell pellet was resuspended in DPBS with 1% Fetal Bovine Serum (FBS). The concentration of BALF immune cells was adjusted to 2000 cells/μl. Samples were submitted to the Advance Analytics Core at the University of North Carolina at Chapel Hill (UNC) School of Medicine for library construction and sequencing.

### Cytokine analyses

Cell-free BALF samples were assayed for various cytokine and chemokine contents using Mouse Cytokine kit (Bio-Rad, Hercules, CA), according to the manufacturer’s recommendations.

### Single-cell suspension preparation and library construction

BALF immune cells were stained with acridine orange and propidium iodide and assessed for viability and concentration using the LUNA-FX7 dual fluorescence cell counter (Logos Biosystems, Annandale, VA) and loaded into a Chromium Next GEM Chip G (10X genomics, Pleasanton, CA) to generate gel-beads in emulsions. Barcoded sequencing libraries were constructed using Chromium Single-cell 3’ Reagent Kit v3.1 (10X genomics, Pleasanton, CA). Single-cell libraries were sequenced using NovaSeq X Plus (Admera Health, South Plainfield, NJ).

### Single-cell RNA sequencing data analyses

Raw FASTQ files were demultiplexed and the unique transcripts with unique molecular identifiers (UMI) were counted using the Cell Ranger pipeline (v8.0.0), and individual reads were aligned to GRCm39 mouse reference genome (10X genomics, Pleasanton, CA). Initial cell-by-gene count matrix was imported into Seurat (v5.1.0) (Hao et al., 2024) using R (v4.4.0; https://www.R-project.org/). Cells with fewer than 800 UMIs, fewer than 500 detected genes, and more than 10% mitochondrial gene content were excluded from downstream analyses. An equal number of cells were randomly selected from each group for the downstream integration process. Data were normalized and scaled using SCTransform function, integrated using IntegrateData function, and clustered with Louvain-Jaccard clustering (resolution = 0.2) using FindNeighbors and FindClusters functions. Uniform manifold approximation and projection (UMAP) analysis was performed using the RunUMAP function (Hao et al., 2024). Cell annotation was performed using a list of predefined markers (**Supplemental Table S2**).

Differential gene expression analyses among different exposure conditions were performed using FindMarkers function (Hao et al., 2024).

### Pathway and function analyses

To determine the biological and functional relevance of differentially-expressed genes (DEGs), we employed ingenuity pathway analysis (IPA; https://www.qiagenbioinformatics.com/products/ingenuity-pathway-analysis) (Krämer et al., 2014). IPA identifies molecular and functional pathways that are influenced by DEGs. DEGs (|average log2FC|= 0.1, adjusted *P* value ≤ 0.05, min.pct = 0.1) were uploaded into IPA software, and their contribution towards the enrichment of canonical pathways was analyzed.

### Construction of the cellular trajectory analysis

R package Monocle3 (v1.3.7; https://cole-trapnell-lab.github.io/monocle3/) (Qiu et al., 2017) was applied to construct the cellular development trajectory along the pseudotime. Working Seurat object was converted to cell dataset (cds) format to execute Monocle3. Cell development trajectory was analyzed using the learn_graph function, and clusters of cells were ordered based on their gene expression along the pseudotime using the order_cells function. Finally, UMAP with the developmental trajectory line was plotted using the plot_cells function.

### Statistical analyses

One-way analysis of variance (ANOVA) followed by the Tukey’s post hoc test for multiple comparisons was used to determine significant differences among groups. All data were expressed as means ± Standard error of the mean (SEM). *P*-value ≤ 0.05 was considered statistically significant. Statistical analyses were performed using GraphPad Prism 8.0.1 (GraphPad Software, La Jolla, CA).

## Results

### Ozone exposure alters immune cell composition and the levels of inflammatory mediators in the airspaces

To determine the concentration-dependent effect of single exposure of ozone (O_3_) on immune cell composition in the lung airspaces, immune cells were quantified in the bronchoalveolar lavage fluid (BALF) from filtered air (FA)-or O_3_-exposed mice (**Figure 1A-C**). While the BALF from FA-exposed mice did not contain neutrophils, the 1 ppm and 1.5 ppm O_3_-exposed mice exhibited significantly increased neutrophil recruitment (**Figure 1B-C**). Although non-significantly, the neutrophil and lymphocyte percentages were higher in 1.5 ppm O_3_-exposed mice versus 1 ppm O_3_-exposed mice (**Figure 1B-C**).

**Figure 1:**
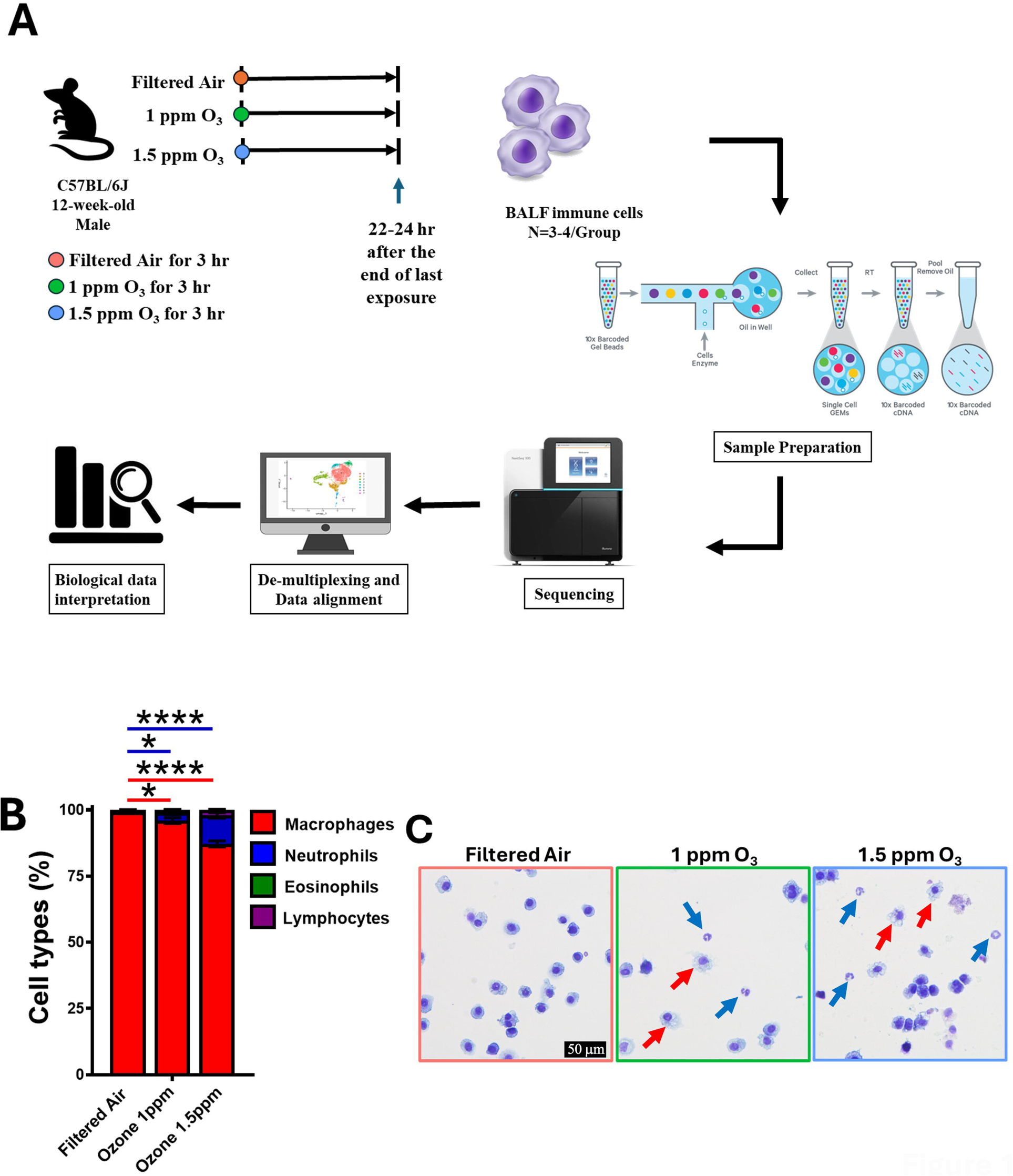
Ozone exposure alters the immune cell composition in the airspaces. **(A)** Schematic diagram depicting the filtered air (FA) and ozone (O_3_) exposure regimen and subsequent scRNA-seq analyses pipeline. **(B)** Percentages of bronchoalveolar lavage (BAL) cell types are shown in a stacked bar graph (macrophages [red], neutrophils [blue], eosinophils [green], and lymphocytes [purple]). Error bars represent standard error of the mean (SEM), \**p* <0.05, \*\*\*\**p* <0.0001 using one-way ANOVA followed by Tukey’s post hoc test for multiple comparisons. **(C)** Representative photomicrographs of Wright-Giemsa-stained BAL cells cytospins from FA-(solid pink border), 1 ppm O_3_-(solid green border), and 1.5 ppm O_3_-exposed mice (solid blue border). Red arrows depict alveolar macrophages. Blue arrows depict neutrophils.

To assess the effect of O_3_ exposure on the release of inflammatory mediators within the lung airspaces, we assessed the levels of selected cytokines and chemokines in the BALF of FA- or O_3_-exposed mice (**Figure 2**). The BALF levels of neutrophil-specific chemokines, i.e., macrophage inflammatory protein 2 (MIP-2/CXCL2), keratinocyte chemoattractant (KC/CXCL1), and LPS-induced CXC chemokine (LIX/CXCL5) trended higher in 1 ppm O_3_-exposed mice versus FA-exposed mice, but the increase was not statistically significant (**Figure 2A-C**). Among the three neutrophil-specific chemokines analyzed, the BALF levels for MIP-2 levels were significantly higher in 1.5 ppm O_3_-exposed mice versus FA-as well as 1 ppm O_3_-exposed mice (**Figure 2A**). The increase in the BALF levels of these neutrophil-specific chemokines was consistent with the increase in neutrophil counts in O_3_-exposed mice as compared with FA-exposed mice.

**Figure 2:**
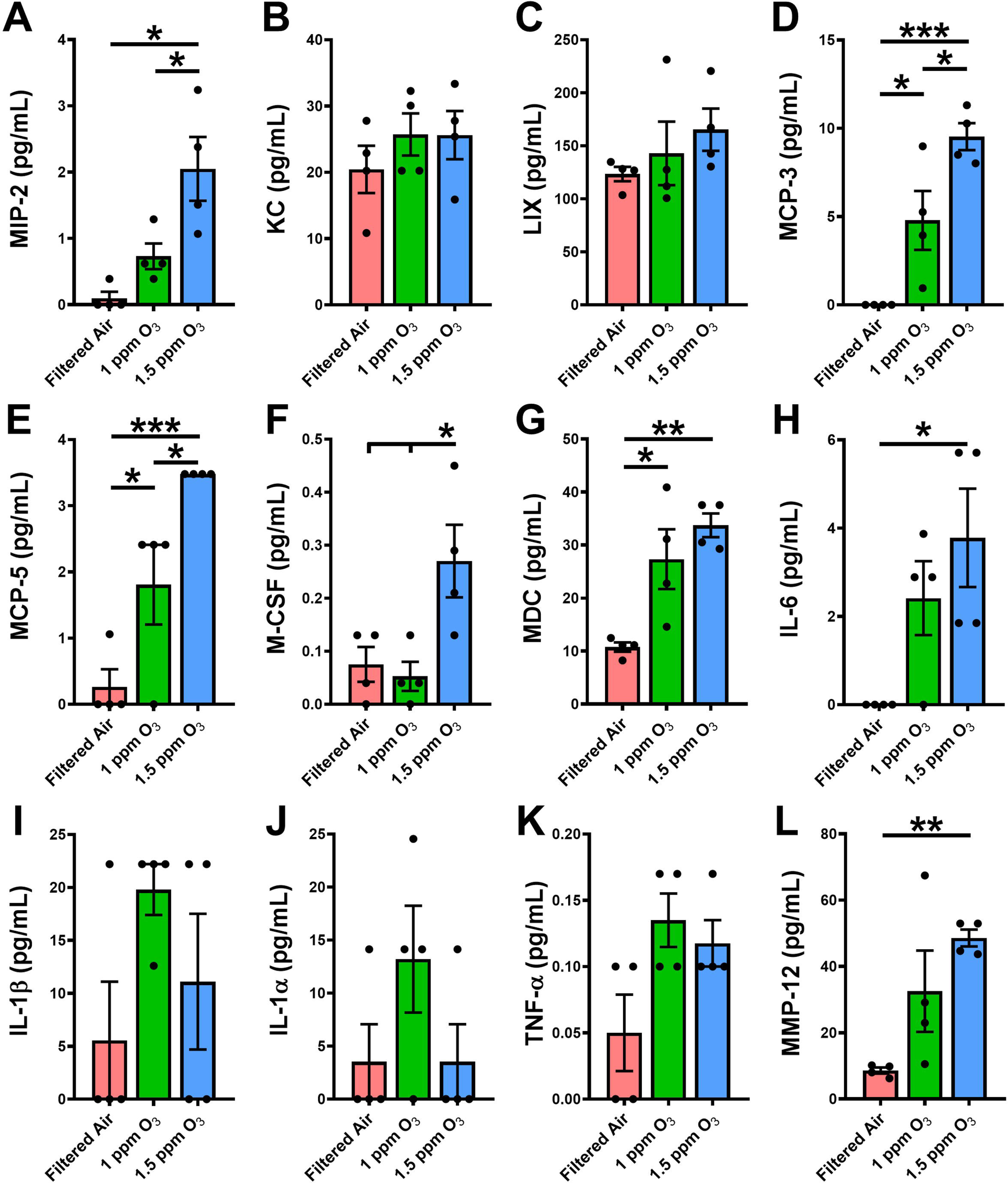
Ozone exposure alters the levels of inflammatory mediators in the airspaces. Cytokine levels (pg/ml; picogram per milliliter) of (**A**) macrophage inflammatory protein 2 (MIP-2/CXCL2), (**B**) keratinocyte chemoattractant (KC/ CXCL1), (**C**) LPS-induced CXC chemokine (LIX/CXCL5), (**D**) monocyte chemotactic protein 3 (MCP-3/CCL7), (**E**) monocyte chemotactic protein 5 (MCP-5/CCL12), (**F**) macrophage colony stimulating factor (M-CSF/CSF1), (**G**) macrophage-derived chemokine (MDC/CCL22), (**H)** interleukin 6 (IL-6), (**I**) interleukin 1 beta (IL-1[3), (**J**) interleukin 1 alpha (IL-1α), (**K**) tumor necrosis factor alpha (TNF-α), (**L**) matrix metallopeptidase 12 (MMP-12) in the BALF from FA-exposed mice (solid pink bar), 1 ppm ozone (O_3_)-exposed mice (solid green bar), and 1.5 ppm O_3_-exposed mice (solid blue bar). Error bars represent standard error of the mean (SEM), \**p*<0.05, \*\**p*<0.01, \*\*\**p*<0.001 using one-way ANOVA followed by Tukey’s post hoc test for multiple comparisons.

Next, we assessed the BALF levels of monocytes-/macrophages-specific cytokines, i.e., monocyte chemotactic protein 3 (MCP-3/CCL7), monocyte chemotactic protein 5 (MCP-5/CCL12), and macrophage colony-stimulating factor (M-CSF/CSF1). MCP-3 and MCP-5 levels were significantly increased in both O_3_-exposed groups as compared with FA-exposed group (**Figure 2D-E**). M-CSF levels were comparable between 1 ppm O_3_-exposed and FA-exposed groups but were significantly increased in 1.5 ppm O_3_-exposed group as compared to 1 ppm O_3_-exposed and FA-exposed groups (**Figure 2F**).

Macrophage-derived chemokine (MDC/CCL22) is responsible for the recruitment of T helper cells type 2 (Th2) and regulatory T cells during lung inflammation (Iellem et al., 2001; Imai et al., 1999). The BALF levels of MDC were significantly elevated in both O_3_-exposed groups as compared with FA-exposed group (**Figure 2G**). MDC levels trended higher in 1.5 ppm O_3_-exposed mice as compared with 1 ppm O_3_-exposed mice, but the increase was not statistically significant (**Figure 2G**).

The levels of various pro-inflammatory cytokines, including interleukin 6 (IL-6), interleukin 1 beta (IL-1[3), interleukin 1 alpha (IL-1α), and tumor necrosis factor alpha (TNF-α) were also assessed. The BALF levels of IL-6 in 1.5 ppm O_3_-exposed mice were significantly elevated as compared with FA-exposed mice and trended higher as compared with 1 ppm O_3_-exposed mice (**Figure 2H**). In contrast, the BALF levels of IL-1α, IL-1[3, and TNF-α trended higher in both O_3_-exposed groups as compared with FA-exposed group (**Figure 2I-K**).

Matrix metallopeptidase 12 (MMP-12) is a macrophage-specific protease that is responsible for the alveolar space enlargement in various disease models, including O_3_-induced lung inflammation (Choudhary, Vo, Paudel, Yadav, et al., 2021; Finlay et al., 1997; Trojanek et al., 2014). MMP-12 levels trended higher in 1 ppm O_3_-exposed mice as compared with FA-exposed mice, but the increase was not statistically significant (**Figure 2L**). The BALF levels of MMP-12 were significantly increased in 1.5 ppm O_3_-exposed mice as compared with FA-exposed mice (**Figure 2L**).

### Ozone exposure results in concentration-dependent transcriptional changes in BALF cells

The BALF cells were collected from FA, 1 ppm, and 1.5 ppm O_3_-exposed groups to generate the single-cell RNA sequencing (scRNA-seq) profiles. An equal number of cells (n = 2213/exposure group) that passed quality control filtering, were randomly selected for further integration steps (**Supplemental Figure S1**). Using a conservative clustering resolution (res = 0.2), the integrated UMAP identified 8 cellular clusters (Cluster 0, 1, 2, 3, 4, 5, 6, 7) (**Figure 3A**). The individual UMAP of BALF cells from three exposure groups showed almost similar pattern of clustering, while differing in the cellular density within the different clusters (**Figure 3B-C**, **Supplemental Table S1**).

**Figure 3:**
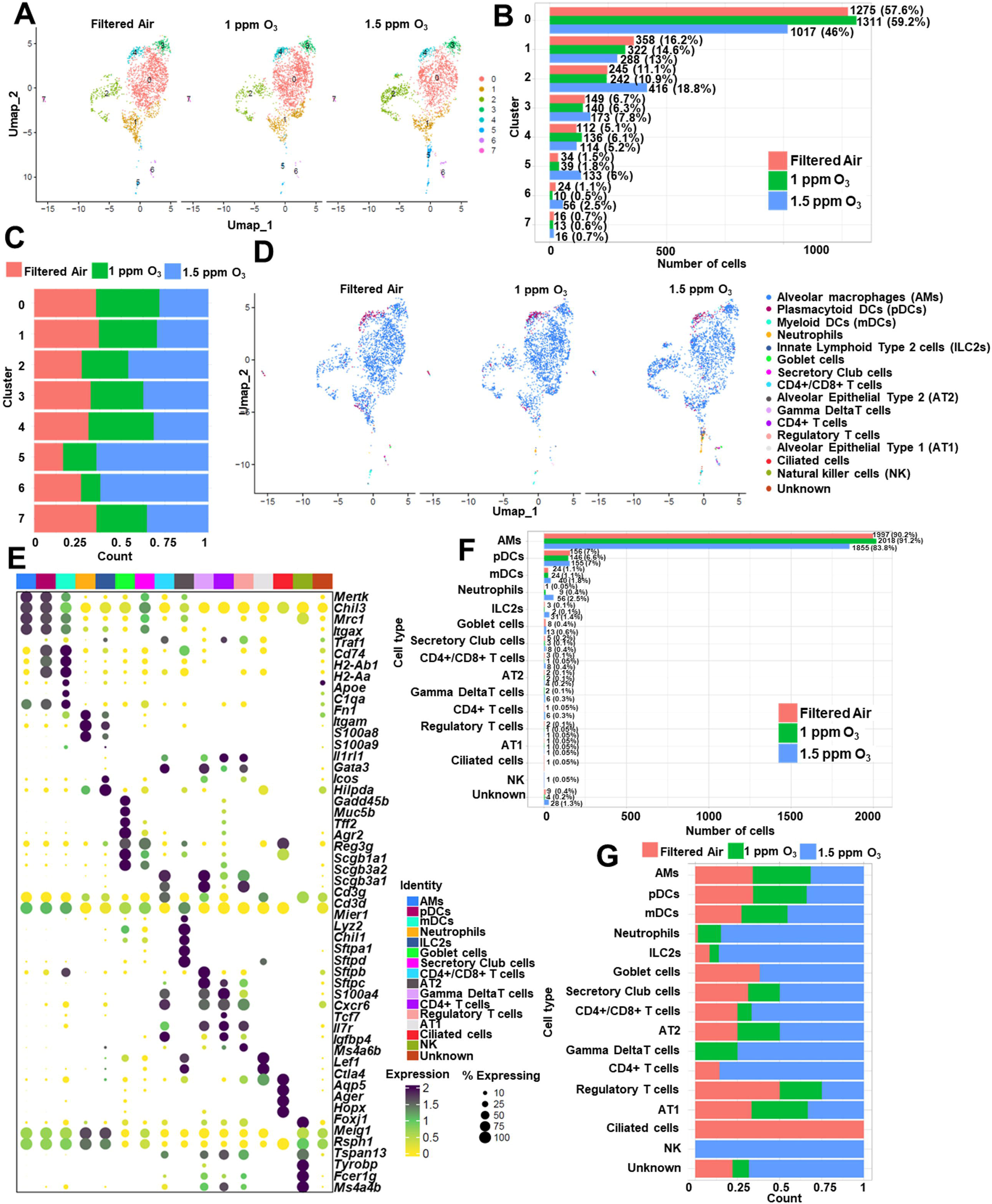
Ozone exposure alters cellular density in the UMAP clusters. **(A)** Uniform manifold approximation and projection (UMAP) plot of scRNA-seq data from filtered air (FA) (n=4), 1 ppm ozone (O_3_) (n=4), and 1.5 ppm O_3_ (n=4) depicting 8 distinct cellular clusters (0-7). **(B)** Bar graph depicting total cell number in each cell cluster from FA (solid pink bar), 1 ppm O_3_ (solid green bar), and 1.5 ppm O_3_ (solid blue bar). **(C)** Stacked bar graph depicting relative cell proportion in each cell cluster from FA (solid pink), 1 ppm O_3_ (solid green), and 1.5 ppm O_3_ (solid blue). **(D)** UMAP plot illustrating major cell populations identified by distinct markers, with each population represented by a different color. **(E)** Dot plot depicting the scaled expression level (yellow (low) to black (high) colored dots) and percentage of cells (size of dots) expressing the designated marker genes for each cell population. **(F)** Bar graph depicting total cell numbers in each cell population from FA (solid pink), 1 ppm O_3_ (solid green), and 1.5 ppm O_3_ (solid blue). **(G)** Stacked bar graph depicting relative cell proportion in each cell population from FA (solid pink), 1 ppm O_3_ (solid green), and 1.5 ppm O_3_ (solid blue).

Cell-specific genes (**Supplemental Table S2**) were used to annotate various cell clusters identified in the UMAP (**Figure 3D-E**). Alveolar macrophages (AMs: *Mertk, Chil3, Lyz2, Mrc1, Itgax*) accounted for the largest proportion of the BALF immune cells from each group [FA = 1997 cells (90.2%); 1 ppm O_3_ = 2018 cells (91.2%); 1.5 ppm O_3_ = 1855 cells (83.8%)] (**Figure 3F-G**). Other predominant cell types in the lung airspaces of FA- and O_3_-exposed mice were plasmacytoid dendritic cells (pDCs; *Traf1, Cd74, H2-Ab1, H2-Aa*) and myeloid dendritic cells (mDCs: *H2-Ab1, H2-Aa, Cd74, Apoe, C1qa, Fn1*). pDCs accounted for the second largest proportion of the BALF immune cells from each group [FA = 156 cells (7%); 1 ppm O_3_ = 146 cells (6.6%); 1.5 ppm O_3_ = 155 cells (7%)] (**Figure 3F-G**). mDCs accounted for the third largest proportion of the BALF immune cells from each group [FA = 24 cells (1.1%); 1 ppm O_3_ = 24 cells (1.1%); 1.5 ppm O_3_ = 40 cells (1.8%)] (**Figure 3F-G**). Additionally, neutrophils (*S100a8, S100a9, Retnlg, Itgam*) and lymphocytes, i.e., CD4+ T cells (*Tcf7, Il7r, Igfbp4, Ms4a6b, Lef1*), CD4+/CD8+ T cells (*Cd3g, Cd3d, Mier1*), type 2 innate lymphoid cells (ILC2s: *Il1rl1, Hilpda, Gadd45b*), were increased in 1.5 ppm O_3_-exposed mice as compared with FA- and 1 ppm O_3_-exposed mice (**Figure 3F-G**).

Alveolar macrophages (AMs) are among the first cellular responders to the inhaled toxicants including O_3_ (Patial & Saini, 2020) and display altered transcriptional and functional responses during O_3_-induced lung injury (Choudhary, Vo, Paudel, Patial, et al., 2021; Choudhary, Vo, Paudel, Wen, et al., 2021; Kumagai et al., 2016, 2017). However, how the transcriptional profile of AMs changes in response to varying concentrations of O_3_ at the single-cell level is yet to be defined. Towards this, we compared expression levels of transcripts in AMs from each group [FA = 1997 cells (90.2%); 1 ppm O_3_ = 2018 cells (91.2%); 1.5 ppm O_3_ = 1855 cells (83.8%)] (**Figure 3F**). Using the cut-off criteria (|average log2FC| ≥ 0.1, adjusted *P* value ≤ 0.05, min.pct ≥ 0.1), the differential gene expression analyses between 1 ppm O_3_ versus FA group identified 1356 differentially-expressed genes (DEGs) (upregulated, 750; downregulated, 606) (**Figure 4A**). Using the cut-off criteria (|average log2FC| ≥ 0.1, adjusted *P* value ≤ 0.05, min.pct ≥ 0.1), the differential gene expression analyses between 1.5 ppm O_3_ versus FA group revealed a relatively higher number of 2980 DEGs (upregulated, 1287; downregulated, 1693) (**Figure 4A**).

**Figure 4:**
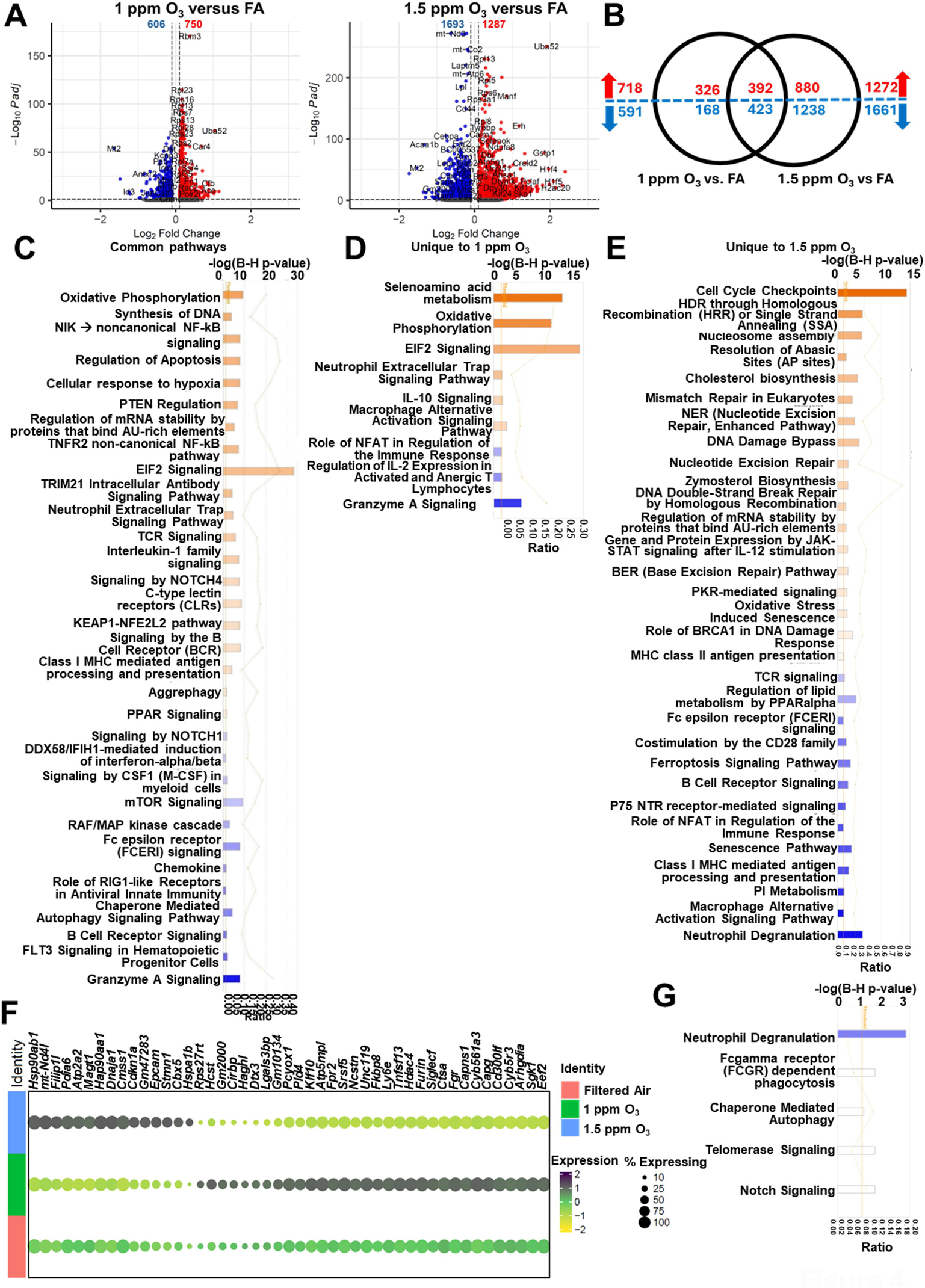
Ozone exposure results in concentration-dependent transcriptomic changes in alveolar macrophages. **(A)** Volcano plots depicting differentially-expressed genes (DEGs; upregulated [red] and downregulated [blue]) (|averageLog2FC | > 0.1; adjusted *P* value ≤ 0.05) in 1 ppm ozone (O_3_) versus filtered air (FA) (left panel) and 1.5 ppm O_3_ versus FA (right panel). **(B)** Venn diagram depicting common and unique DEGs (upregulated [red] and downregulated [blue]) in 1 ppm O_3_ versus FA and 1.5 ppm O_3_ versus FA. **(C)** Bar graph depicting commonly enriched biological pathways between 1 ppm O_3_ and 1.5 ppm O_3_ with respect to FA identified using ingenuity pathway analyses (IPA) approach. Pathway activation (orange) and suppression (blue) status were determined based on IPA z-score. **(D)** Bar graph depicting uniquely enriched biological pathways in 1 ppm O_3_ versus FA identified using IPA approach. Pathway activation (orange) and suppression (blue) status were determined based on IPA z-score. **(E)** Bar graph depicting uniquely enriched biological pathways in 1.5 ppm O_3_ versus FA identified using IPA approach. Pathway activation (orange) and suppression (blue) status were determined based on IPA z-score. **(F)** Dot plot depicting the scaled expression level (yellow (low) to black (high) colored dots) and percentage of cells (size of dots) for genes with expression in opposite direction between 1 ppm O_3_ and 1.5 ppm O_3_ with respect to FA. **(G)** Bar graph depicting enriched biological pathways in 1.5 ppm O_3_ versus 1 ppm O_3_ identified using IPA approach. Pathway suppression (blue) status was determined based on IPA z-score. White color bar indicates z-score=0.

Among the DEGs identified by differential expression analyses, a total of 815 DEGs (upregulated, 392; downregulated, 423) were common to O_3_ exposure, regardless of O_3_ concentration, e.g., *S100a4, Ffar2, Cfb, Car4, S100a6, Gstp1, Fdps, Sqle, Cd14, Tmem223, Fn1, Mt2,* and *Il17d* (**Figure 4B**, **Table 1, Supplemental Table S3**). Additionally, we identified 494 DEGs (upregulated, 326; downregulated, 168) that were differentially expressed exclusively in 1 ppm O_3_ group versus FA group, e.g., *Apoc1, Ace, Scd2, P2ry13,* and *S100a8* (**Figure 4B**, **Table 1, Supplemental Table S3**). On the other hand, we identified 2118 DEGs (upregulated, 880; downregulated, 1238) that were differentially expressed exclusively in 1.5 ppm O_3_ group versus FA group, e.g., *Il1b, Osm, Ccr5, Mki67, Bpifa1, Scgb1a1, Ctsf, Ccl9, Slc27a1, Csf3r*, and *Cd69* (**Figure 4B**, **Table 1, Supplemental Table S3**). The complete list of DEGs is included in **Supplemental Table S3**.

**Table 1.**
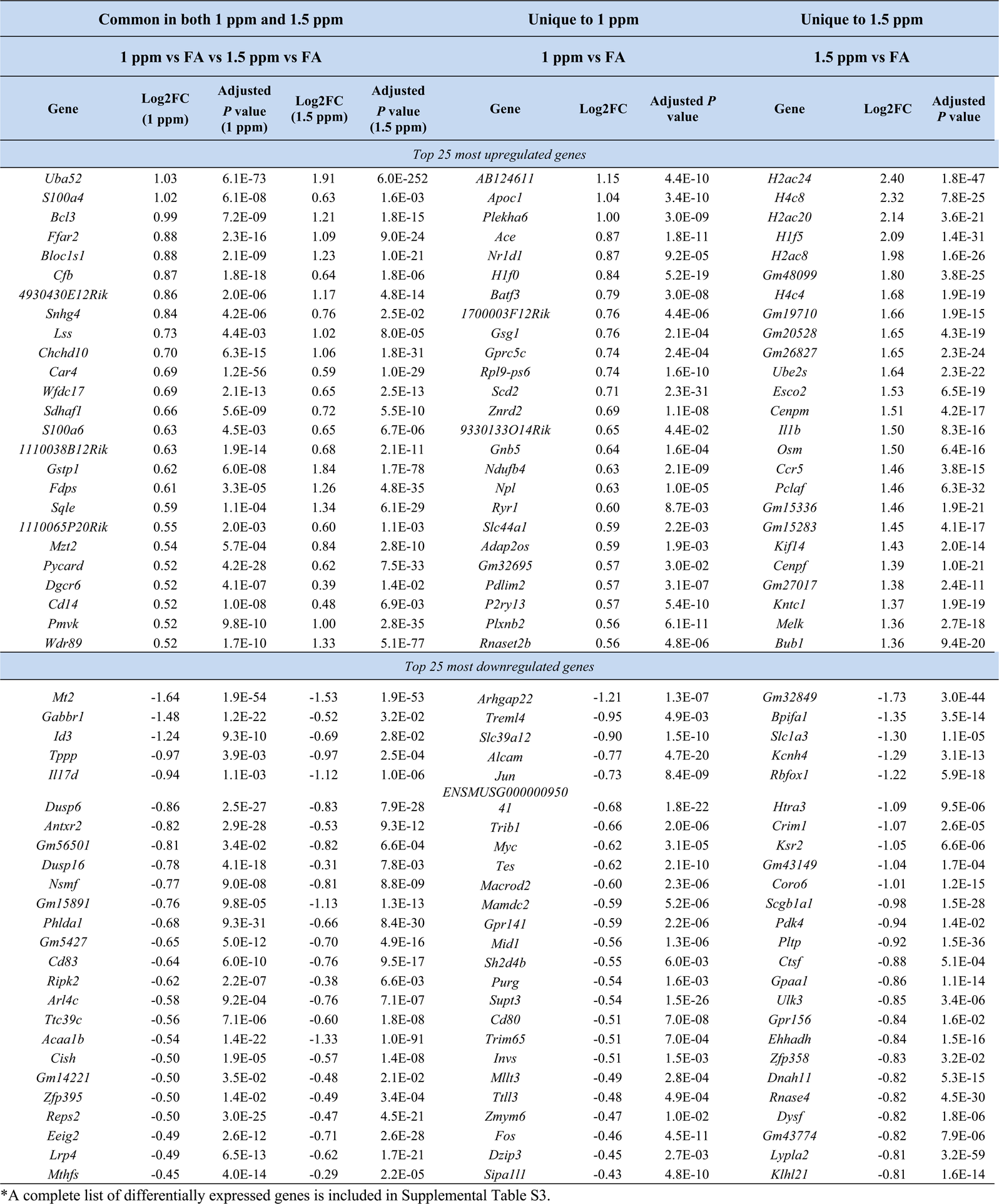
Top 25 most upregulated and top 25 most downregulated genes in AMs from 1 ppm O_3_ and 1.5 ppm O_3_ (common and unique) with respect to filtered air (FA).

Ingenuity pathway analysis (IPA) revealed that common DEGs identified in 1 ppm O_3_ versus FA group and 1.5 ppm O_3_ versus FA group comparisons suggested the enrichment of pathways, including oxidative phosphorylation, non-canonical NF-kB signaling, regulation of apoptosis, cellular response to hypoxia, regulation of mRNA stability by proteins that bind AU-rich elements, TRIM21 intracellular antibody signaling, neutrophil extracellular trap signaling, interleukin-1 family signaling, NOTCH4 signaling, and the suppression of pathways including mTOR signaling, chemokine signaling, and Fc epsilon receptor (FCERI) signaling (**Figure 4C, Supplemental Table S4**).

The unique DEGs from 1 ppm O_3_ versus FA comparison were associated with the enrichment of pathways, including selenoamino acid metabolism, IL-10 signaling, and macrophage alternative activation signaling (**Figure 4D, Supplemental Table S4**). Meanwhile, unique DEGs from 1.5 ppm O_3_ versus FA comparison were associated with the enrichment of pathways including cell cycle checkpoints, DNA damage bypass and repair, cholesterol biosynthesis, oxidative stress-induced senescence, MHC class II antigen presentation, and the suppression of pathways including regulation of lipid metabolism by PPARalpha, ferroptosis signaling, role of NFAT in regulation of the immune response, senescence pathway, and macrophage alternative activation signaling pathway (**Figure 4E, Supplemental Table S4**).

Differential expression analyses also revealed genes that were differentially expressed in opposite direction in 1 ppm O_3_ group versus FA group and 1.5 ppm O_3_ group versus FA group. We identified 15 DEGs that were upregulated in 1.5 ppm O_3_ versus FA but were downregulated in 1 ppm O_3_ versus FA, e.g., *Filip1l, Pdia6, Cdkn1a, Stmn1, Cbx5* and *Hspa1b* (**Figure 4F, Supplemental Table S3**) On the other hand, we identified 32 DEGs that were downregulated in 1.5 ppm O_3_ versus FA but were upregulated in 1 ppm O_3_ versus FA, e.g., *Klf10, Tnfsf13, Ctsa, Fgr, Capns1, Cd300lf,* and *Sgk1* (**Figure 4F, Supplemental Table S3**). IPA revealed that as compared to 1 ppm O_3_ concentration, exposure to 1.5 ppm O_3_ suppressed neutrophil degranulation. Other pathways that were enriched in 1.5 ppm O_3_ mice versus 1 ppm O_3_ mice include Fc gamma receptor (FCGR) dependent phagocytosis, chaperone mediated autophagy, telomerase signaling, and notch signaling (**Figure 4G**).

### AM heterogeneity post-ozone exposure

Next, we generated UMAP subclusters based on the positive expression for all the selected AM-specific signatures, i.e., *Mertk, Chil3, Lyz2, Mrc1, Itgax*, which resulted in the 5 subclusters, i.e., AM clusters 0, 1, 2, 3, and 4 (**Figure 5A**). AM cluster 0 [FA = 818 cells (35.8%); 1 ppm O_3_ = 866 cells (37.9%); 1.5 ppm O_3_ = 601 cells (26.3%)] (**Figure 5B-C, Supplemental Table S5**) was defined by the expression of markers, including *Cd44, Ahnak*, *Afdn, Mmp19,* and *Rasa2* (**Figure 5D, Supplemental Table S6)**. AM cluster 1 [FA = 457 cells (34%); 1 ppm O_3_ = 476 cells (35.3%); 1.5 ppm O_3_ = 414 cells (30.7%)] was defined by the expression of markers, including *Rpl35a, Rps27a, Rps14, Rps13,* and *Ear2* (**Figure 5D, Supplemental Table S6**). AM cluster 2 [FA = 344 cells (36.7%); 1 ppm O_3_ = 305 cells (32.6%); 1.5 ppm O_3_ = 288 cells (30.7%)] was defined by the expression of markers, including *Sqstm1, Cat, Afp, Acod1,* and *Efr3b* (**Figure 5D, Supplemental Table S6**). AM cluster 3 [FA = 240 cells (27.3%); 1 ppm O_3_ = 240 cells (27.3%); 1.5 ppm O_3_ = 399 cells (45.4%)] was defined by the expression of markers, including *Tmpo, H2az2, Pclaf, Cep55,* and *Cdca3* (**Figure 5D, Supplemental Table S6**). Finally, AM cluster 4 [FA = 138 cells (32.7%); 1 ppm O_3_ = 131 cells (31%); 1.5 ppm O_3_ = 153 cells (36.3%)] was defined by the expression of markers, including *Malat1, Gab2, Macf1, Ssh2,* and *Gm27008* (**Figure 5D, Supplemental Table S6**).

**Figure 5:**
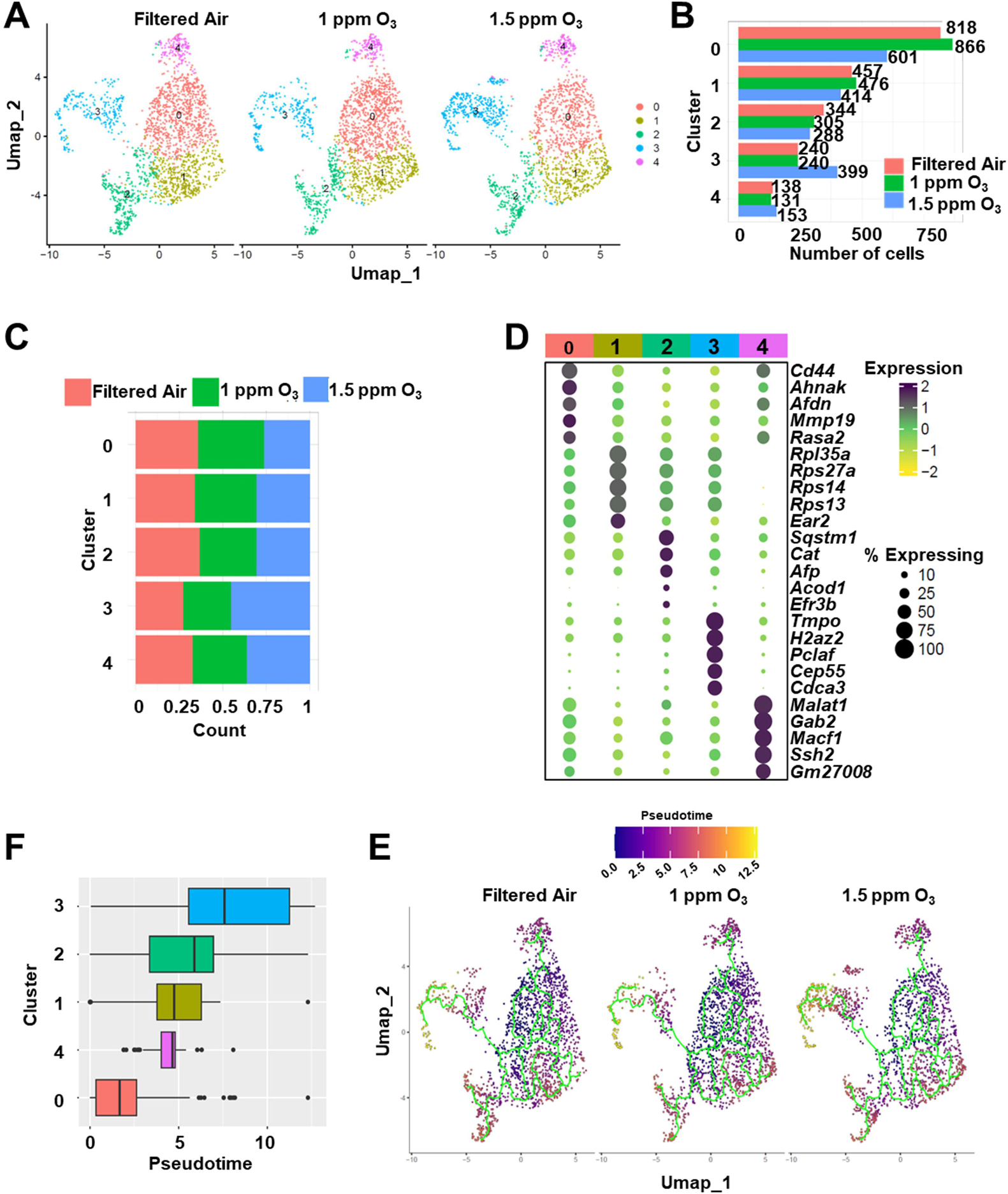
AM displays heterogeneity post-ozone exposure. **(A)** UMAP plot of annotated AMs from filtered air (FA), 1 ppm ozone (O_3_), and 1.5 ppm O_3_ depicting five distinct AM subclusters (0-4). **(B)** Bar graph depicting total AM counts in each cluster from FA (solid pink bar), 1 ppm O_3_ (solid green bar), and 1.5 ppm O_3_ (solid blue bar). **(C)** Stacked bar graph depicting relative cell proportion in each AM cluster from FA (solid pink color), 1 ppm O_3_ (solid green color), and 1.5 ppm O_3_ (solid blue color). **(D)** Dot plot depicting the scaled expression level (color) and percent expressing (size) for defined marker genes for each AM subcluster. **(E)** UMAP plot colored by pseudotime representing differentiation from dark blue (less differentiated) to yellow (more differentiated) in FA (left), 1 ppm O_3_ (middle), 1.5 ppm O_3_ (right). **(F)** Boxplots of the range of pseudotime in each subcluster, colored by subcluster annotation.

AM is a heterogenous population (Dick et al., 2022; Wu et al., 2023; Xu-Vanpala et al., 2020). First, to determine whether the different AM clusters represent cell populations at different stages of development, we performed Trajectory analyses (Qiu et al., 2017), which assess gene expression changes to determine the cellular differentiation stage of each cluster. Trajectory analysis revealed that the differentiation of AMs began with the most inactive cluster (AM cluster 0) and progressed through AM clusters 4, 1, 2 and ultimately to cluster 3. However, interestingly, the trajectories of AM clusters between the three treatment groups remained unaltered (**Figure 5E-F**).

### Ozone exposure alters transcriptome of AMs in a concentration- and cluster-dependent manner

Comparison of AM gene expression between different treatment groups revealed several differences in a cluster-specific manner. AM cluster 0, which accounts for ∼39% (2285 out of 5870; **Supplemental Table S5**) of total annotated AMs, was analyzed to identify DEGs between 1 ppm O_3_ versus FA group and 1.5 ppm O_3_ versus FA group. Using the cut-off criteria (|average log2FC| ≥ 0.1, adjusted *P* value ≤ 0.05, min.pct ≥ 0.1), the analyses identified 712 DEGs (upregulated, 343; downregulated, 369) in 1 ppm O_3_ versus FA, and 1357 DEGs (upregulated, 551; downregulated, 806) in 1.5 ppm O_3_ versus FA group (**Figure 6A, Supplemental Table S7)**. The comparison of 2069 (712+1357) identified DEGs revealed a total of 401 DEGs (upregulated, 185; downregulated, 216) that were common to 1 ppm O_3_ and 1.5 ppm O_3_ groups with respect to FA group (**Figure 6B**, **Table 2, Supplemental Table S7**). The IP analyses on these 401 DEGs suggested that O_3_ exposure to either concentration caused enrichment of pathways involved in active translation responses (**Figure 7A, Supplemental Table S12**). We identified 293 DEGs (upregulated, 149; downregulated, 144) that were uniquely expressed in 1 ppm O_3_ versus FA group (**Figure 6B**, **Table 2, Supplemental Table S7**). The IP analyses on these 293 DEGs suggested enrichment of pathways involved in active translation responses, mitochondrial electron transport chain complex assembly and biogenesis, IL-10 signaling, and PPAR signaling (**Figure 7A, Supplemental Table S12**). Finally, 938 DEGs (upregulated, 357; downregulated, 581) were uniquely expressed in 1.5 ppm O_3_ versus FA group (**Figure 6B**, **Table 2, Supplemental Table S7**). The IP analyses on these 938 DEGs suggested enrichment of pathways involved in cholesterol biosynthesis (**Figure 7A, Supplemental Table S12**).

**Figure 6:**
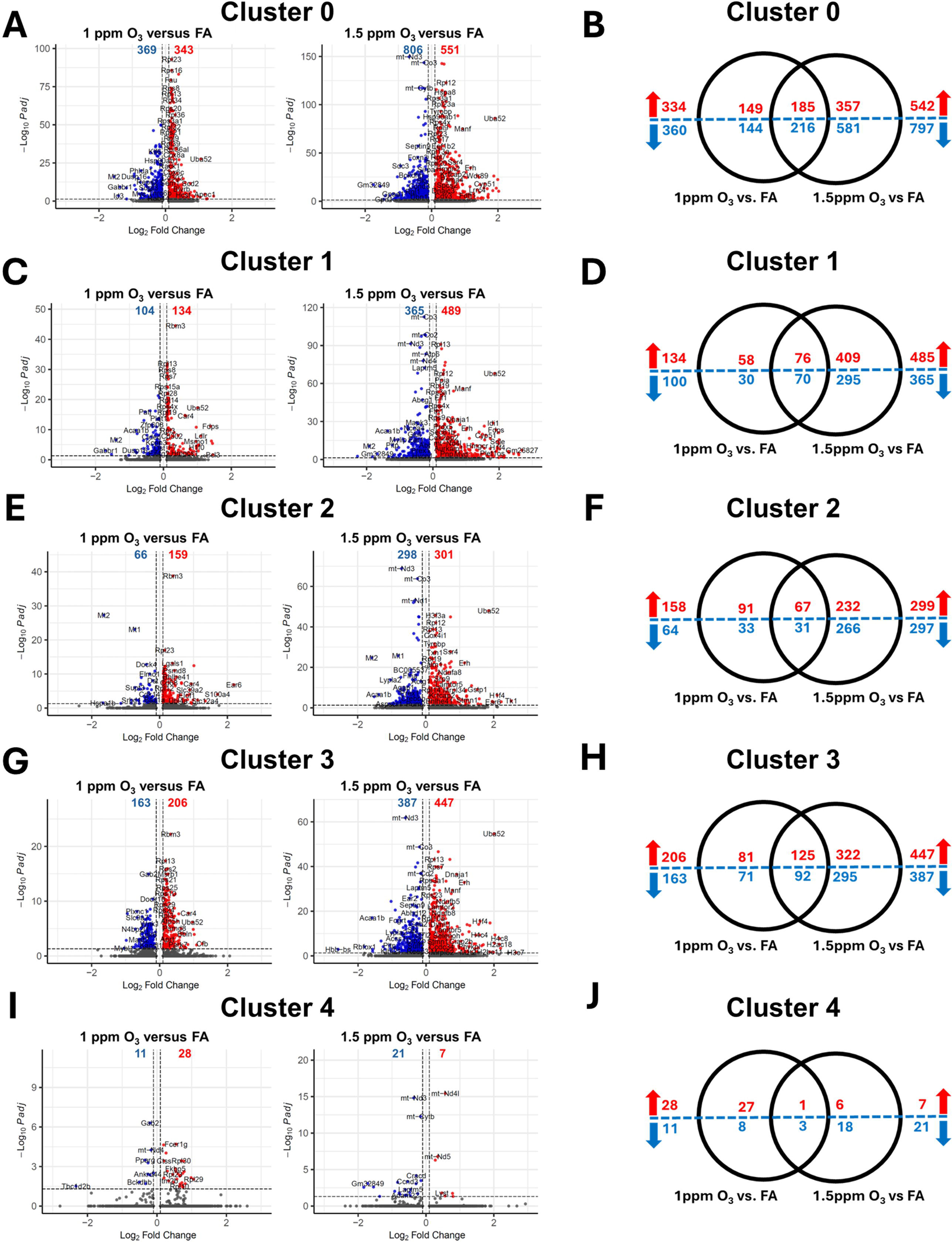
Ozone exposure alters the transcriptome of AMs in a concentration- and cluster-dependent manner. **(A)** Volcano plots depicting differentially-expressed genes (DEGs; upregulated [red] and downregulated [blue]) in AM cluster 0 (|averageLog2FC | > 0.1; adjusted *P* value ≤ 0.05) in 1 ppm ozone (O_3_) versus FA (left panel) and 1.5 ppm O_3_ versus FA (right panel). **(B)** Venn diagram depicting common and unique DEGs in AM cluster 0 (upregulated [red] and downregulated [blue]) in 1 ppm O_3_ versus FA and 1.5 ppm O_3_ versus FA. **(C)** Volcano plots depicting DEGs (upregulated [red] and downregulated [blue]) in AM cluster 1 (|averageLog2FC | > 0.1; adjusted *P* value ≤ 0.05) in 1 ppm ozone (O_3_) versus FA (left) and 1.5 ppm O_3_ versus FA (right). **(D)** Venn diagram depicting common and unique DEGs in AM cluster 1 (upregulated [red] and downregulated [blue]) in 1 ppm O_3_ versus FA and 1.5 ppm O_3_ versus FA. **(E)** Volcano plots depicting DEGs (upregulated [red] and downregulated [blue]) in AM cluster 2 (|averageLog2FC | > 0.1; adjusted *P* value ≤ 0.05) in 1 ppm ozone (O_3_) versus FA (left) and 1.5 ppm O_3_ versus FA (right). **(F)** Venn diagram depicting common and unique DEGs in AM cluster 2 (upregulated [red] and downregulated [blue]) in 1 ppm O_3_ versus FA and 1.5 ppm O_3_ versus FA. **(G)** Volcano plots depicting DEGs (upregulated [red] and downregulated [blue]) in AM cluster 3 (|averageLog2FC | > 0.1; adjusted *P* value ≤ 0.05) in 1 ppm ozone (O_3_) versus FA (left) and 1.5 ppm O_3_ versus FA (right). **(H)** Venn diagram depicting common and unique DEGs in AM cluster 3 (upregulated [red] and downregulated [blue]) in 1 ppm O_3_ versus FA and 1.5 ppm O_3_ versus FA. **(I)** Volcano plots depicting DEGs (upregulated [red] and downregulated [blue]) in AM cluster 4 (|averageLog2FC | > 0.1; adjusted *P* value ≤ 0.05) in 1 ppm ozone (O_3_) versus FA (left) and 1.5 ppm O_3_ versus FA (right). **(J)** Venn diagram depicting common and unique DEGs in AM cluster 4 (upregulated [red] and downregulated [blue]) in 1 ppm O_3_ versus FA and 1.5 ppm O_3_ versus FA.

**Figure 7:**
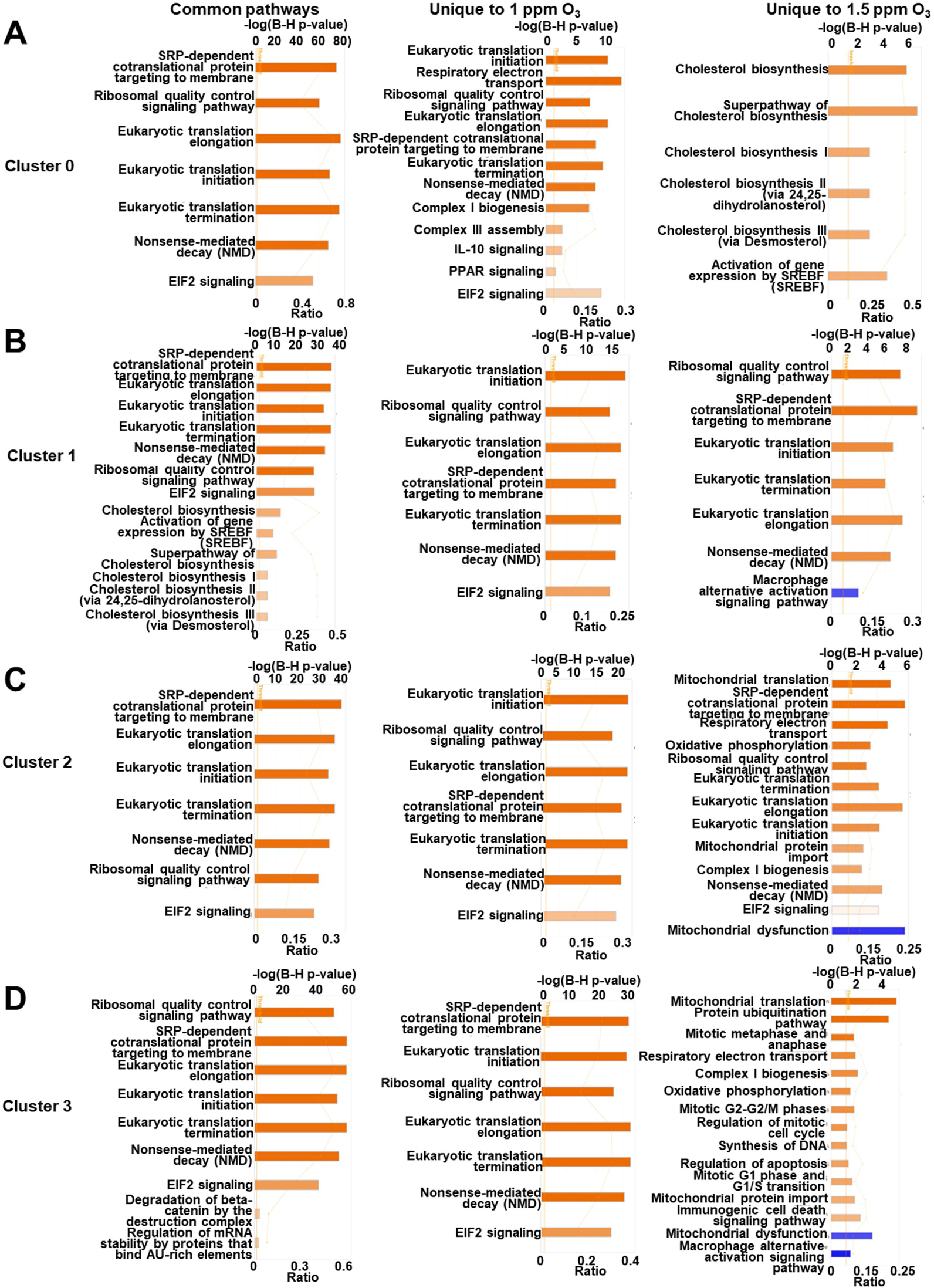
Ozone exposure alters the biological pathways of AMs in a concentration- and cluster-dependent manner. Bar graphs depicting commonly (left panels), uniquely enriched biological pathway in 1 ppm O_3_ versus FA (middle panels), and uniquely enriched biological pathways in 1.5 ppm O_3_ versus FA (right panels) in (**A**) AM cluster 0, (**B**) AM cluster 1, (**C**) AM cluster 2, and (**D**) AM cluster 3 identified using ingenuity pathway analyses (IPA) approach Pathway activation (orange) and suppression (blue) status were determined based on IPA z-score.

**Table 2.**
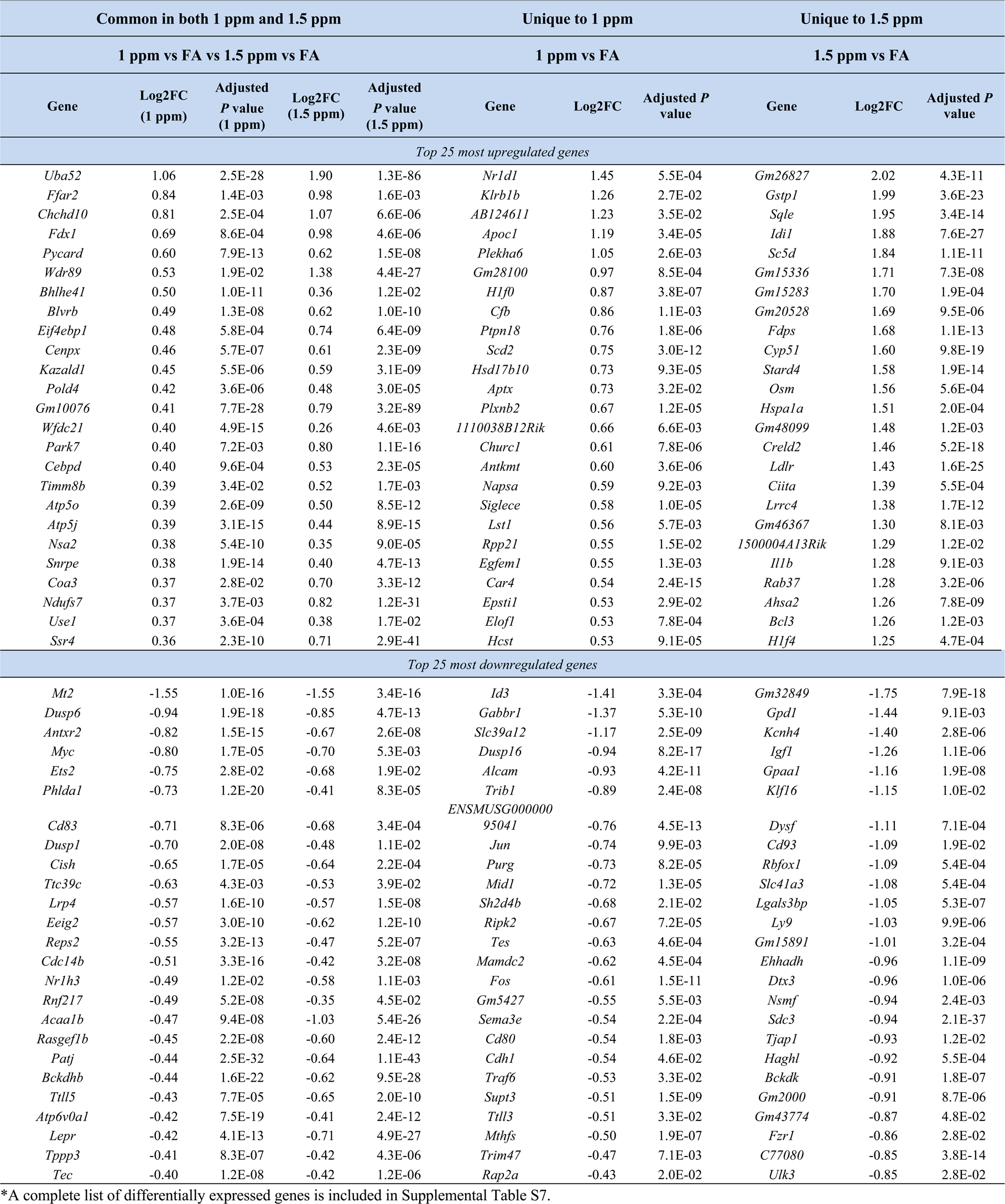
Top 25 most upregulated and top 25 most downregulated genes in Cluster 0 AMs from 1 ppm O_3_ and 1.5 ppm O_3_ (common and unique) with respect to filtered air (FA).

AM cluster 1, which accounts for ∼23% (1347 out of 5870; **Supplemental Table S5**) of total annotated AMs, was analyzed to identify DEGs between 1 ppm O_3_ versus FA and 1.5 ppm O_3_ versus FA group. Using the cut-off criteria (|average log2FC| ≥ 0.1, adjusted *P* value ≤ 0.05, min.pct ≥ 0.1), our analyses identified 238 DEGs (upregulated, 134; downregulated, 104) in 1 ppm O_3_ versus FA, and 854 DEGs (upregulated, 489; downregulated, 365) in 1.5 ppm O_3_ versus FA (**Figure 6C, Supplemental Table S8**). The comparison of 1092 (238+854) identified DEGs revealed a total of 146 DEGs (upregulated, 76; downregulated, 70) that were common to 1 ppm O_3_ and 1.5 ppm O_3_ with respect to FA-exposed mice (**Figure 6D**, **Table 3, Supplemental Table S8**). The IP analyses on these 146 DEGs revealed that the O_3_ exposure to either concentration causes enrichment of pathways involved in active translation responses and cholesterol biosynthesis (**Figure 7B, Supplemental Table S12**). We identified 88 DEGs (upregulated, 58; downregulated, 30) that were uniquely expressed in 1 ppm O_3_ versus FA group (**Figure 6D**, **Table 3, Supplemental Table S8**). The IP analyses on these 88 DEGs suggested an enrichment of protein translation pathways (**Figure 7B, Supplemental Table S12**). Finally, 704 DEGs (upregulated, 409; downregulated, 295) were uniquely expressed in 1.5 ppm O_3_ versus FA group (**Figure 6D**, **Table 3, Supplemental Table S8**). The IP analyses on these 704 DEGs suggested enrichment of active translation responses and suppression of alternate activation of macrophages (**Figure 7B, Supplemental Table S12**).

**Table 3.** Top 25 most upregulated and top 25 most downregulated genes in Cluster 1 AMs from 1 ppm O_3_ and 1.5 ppm O_3_ (common and unique) with respect to filtered air (FA).

| Common in both 1 ppm and 1.5 ppm |  |  |  |  | Unique to 1 ppm |  |  | Unique to 1.5 ppm |  |  |
| --- | --- | --- | --- | --- | --- | --- | --- | --- | --- | --- |
| 1 ppm vs FA vs 1.5 ppm vs FA |  |  |  |  | 1 ppm vs FA |  |  | 1.5 ppm vs FA |  |  |
| Gene | Log2FC<br>(1 ppm) | Adjusted<br>P value<br>(1 ppm) | Log2FC<br>(1.5 ppm) | Adjusted<br>P value<br>(1.5 ppm) | Gene | Log2FC | Adjusted P<br>value | Gene | Log2FC | Adjusted P<br>value |
| Top 25 most upregulated genes |  |  |  |  |  |  |  |  |  |  |
| *Bcl3* | 1.46 | 2.2E-02 | 1.79 | 1.1E-03 | *Tlr13* | 1.03 | 1.26E-03 | *H2ac24* | 3.56 | 6.0E-08 |
| *Sc5d* | 1.45 | 3.9E-06 | 1.99 | 2.1E-13 | *Hlf0* | 0.98 | 7.19E-05 | *Gm26827* | 2.69 | 4.9E-08 |
| *Sqle* | 1.41 | 6.3E-07 | 1.96 | 8.3E-15 | *Rpl9-ps6* | 0.91 | 1.79E-03 | *Gm20528* | 2.60 | 2.1E-04 |
| *Fdps* | 1.39 | 7.6E-12 | 1.91 | 4.6E-24 | *Hlf2* | 0.84 | 1.33E-03 | *Ccr5* | 2.58 | 1.4E-05 |
| *Ldlr* | 1.10 | 1.2E-08 | 1.55 | 4.4E-27 | *Sdhaf1* | 0.84 | 1.03E-02 | *6530409C15Rik* | 2.44 | 2.9E-05 |
| *Erg28* | 1.05 | 1.2E-05 | 1.49 | 4.3E-11 | *Tubb6* | 0.79 | 2.73E-04 | *Ciita* | 2.30 | 3.6E-06 |
| *Fdft1* | 1.02 | 1.7E-05 | 1.56 | 3.4E-16 | *2300009A05Rik* | 0.78 | 1.69E-02 | *Insyn2b* | 2.28 | 3.8E-07 |
| *Uba52* | 1.01 | 5.4E-18 | 1.86 | 1.1E-68 | *P2ry13* | 0.76 | 1.01E-02 | *Gm16599* | 2.16 | 1.4E-03 |
| *Cyp51* | 0.97 | 1.9E-04 | 1.60 | 7.5E-20 | *Fkbp5* | 0.63 | 1.79E-15 | *Pclaf* | 2.15 | 2.2E-02 |
| *Idi1* | 0.97 | 2.9E-05 | 1.82 | 1.7E-29 | *Hcst* | 0.59 | 2.55E-04 | *Acss2* | 2.13 | 1.2E-02 |
| *Scd2* | 0.97 | 1.6E-11 | 0.80 | 3.4E-04 | *Polr2e* | 0.52 | 1.47E-03 | *Socs3* | 2.07 | 2.0E-03 |
| *Msmo1* | 0.96 | 8.2E-07 | 1.21 | 5.4E-13 | *Bhlhe41* | 0.49 | 4.62E-05 | *Osm* | 2.02 | 4.9E-09 |
| *Stard4* | 0.94 | 1.8E-02 | 1.73 | 2.4E-17 | *Tubb4b* | 0.47 | 9.65E-03 | *Gm27008* | 2.00 | 2.5E-16 |
| *Ffar2* | 0.85 | 1.6E-02 | 1.31 | 2.6E-08 | *Cd63* | 0.47 | 3.64E-02 | *Gm15764* | 1.99 | 2.9E-02 |
| *Insig1* | 0.81 | 1.1E-02 | 1.11 | 4.1E-08 | *Scarb1* | 0.43 | 2.08E-02 | *Gstp1* | 1.99 | 6.7E-22 |
| *Pmvk* | 0.80 | 5.2E-04 | 1.44 | 1.7E-18 | *Tecr* | 0.42 | 4.20E-02 | *Hlf4* | 1.96 | 2.9E-10 |
| *Wdr89* | 0.74 | 3.4E-03 | 1.50 | 1.1E-20 | *Gas5* | 0.40 | 7.67E-03 | *Gm48678* | 1.94 | 1.1E-09 |
| *Rras* | 0.69 | 2.3E-02 | 0.92 | 8.4E-05 | *Ndufa11* | 0.36 | 1.34E-02 | *Il1b* | 1.92 | 1.2E-07 |
| *Mrpl22* | 0.69 | 9.4E-03 | 0.75 | 2.4E-02 | *Rbm3* | 0.36 | 3.08E-45 | *Gm16337* | 1.92 | 4.2E-07 |
| *Car4* | 0.66 | 2.7E-15 | 0.76 | 2.4E-20 | *Ctisk* | 0.34 | 1.09E-02 | *Gm15283* | 1.85 | 6.2E-03 |
| *Hmgcs1* | 0.63 | 5.7E-03 | 1.10 | 2.1E-17 | *Psmel* | 0.34 | 2.46E-02 | *A330040F15Rik* | 1.81 | 2.2E-04 |
| *Sreb2* | 0.58 | 3.8E-03 | 0.81 | 2.3E-05 | *Lilrb4a* | 0.31 | 2.05E-03 | *Plet1os* | 1.80 | 1.3E-05 |
| *Pycard* | 0.56 | 3.8E-06 | 0.49 | 1.9E-02 | *Tmem258* | 0.30 | 5.16E-03 | *Lysmd4* | 1.78 | 1.7E-06 |
| *Blvrb* | 0.45 | 5.3E-04 | 0.57 | 2.0E-08 | *Atpif1* | 0.29 | 9.70E-06 | *Gm15336* | 1.77 | 4.1E-04 |
| *Wfdc21* | 0.44 | 8.9E-09 | 0.41 | 1.8E-07 | *Srsf5* | 0.28 | 1.19E-03 | *Lrrc4* | 1.73 | 2.9E-09 |
| Top 25 most downregulated genes |  |  |  |  |  |  |  |  |  |  |
| *Mt2* | -1.39 | 2.8E-07 | -1.92 | 2.9E-11 | *Gabbr1* | -1.70 | 8.72E-04 | *Scgb3a1* | -2.28 | 2.7E-03 |
| *Acaa1b* | -0.80 | 1.7E-10 | -1.37 | 2.2E-23 | *Id3* | -1.53 | 1.63E-02 | *Bpifa1* | -1.83 | 7.2E-03 |
| *Dusp6* | -0.79 | 4.3E-03 | -0.82 | 1.6E-03 | *Dusp16* | -0.84 | 6.94E-04 | *Gm32849* | -1.68 | 3.3E-05 |
| *Phlda1* | -0.77 | 2.2E-09 | -0.68 | 4.7E-06 | *Antxr2* | -0.81 | 2.95E-03 | *Scgb1a1* | -1.40 | 3.9E-09 |
| *Tppp3* | -0.58 | 1.9E-04 | -0.68 | 3.1E-06 | *Herpud1* | -0.59 | 4.21E-07 | *Rbfox1* | -1.33 | 5.7E-03 |
| *Arl13b* | -0.58 | 3.7E-03 | -0.74 | 9.5E-06 | *Afdn* | -0.49 | 1.11E-02 | *Cirbp* | -1.23 | 2.4E-02 |
| *Lepr* | -0.57 | 8.8E-14 | -0.83 | 2.4E-22 | *Csnk1e* | -0.45 | 1.15E-02 | *Pltp* | -1.22 | 4.7E-12 |
| *Patj* | -0.54 | 1.3E-16 | -0.76 | 5.4E-27 | *Tec* | -0.44 | 6.01E-03 | *Dtx3* | -1.19 | 2.0E-03 |
| *Bckdhh* | -0.49 | 7.4E-09 | -0.64 | 3.6E-13 | *Dse* | -0.39 | 1.19E-02 | *Acot2* | -1.15 | 1.6E-04 |
| *Plxnc1* | -0.47 | 1.2E-07 | -0.57 | 5.7E-09 | *Ptpre* | -0.32 | 2.06E-03 | *Abca1* | -1.15 | 4.2E-16 |
| *Il1rn* | -0.47 | 3.6E-05 | -0.45 | 1.9E-03 | *Cmss1* | -0.28 | 5.98E-04 | *Dnah11* | -1.11 | 3.1E-02 |
| *Adam22* | -0.45 | 1.6E-02 | -0.63 | 3.2E-06 | *Dennd4a* | -0.27 | 1.86E-03 | *Sgms2* | -1.09 | 2.5E-08 |
| *Slc9a7* | -0.45 | 2.6E-04 | -0.63 | 1.3E-07 | *Stk17b* | -0.24 | 8.05E-09 | *Mylip* | -1.05 | 2.0E-16 |
| *Peli2* | -0.44 | 3.4E-02 | -0.49 | 8.7E-03 | *AI504432* | -0.22 | 2.73E-04 | *Gm5427* | -1.03 | 4.2E-04 |
| *Lpin1* | -0.39 | 5.7E-17 | -0.51 | 5.1E-22 | *Jund* | -0.21 | 5.06E-06 | *Spns1* | -1.03 | 3.9E-10 |
| *Tob1* | -0.36 | 7.1E-03 | -0.40 | 3.1E-03 | *Fndc3a* | -0.21 | 4.96E-02 | *Mepce* | -1.02 | 9.6E-03 |
| *Cidec* | -0.36 | 2.7E-08 | -0.58 | 1.6E-18 | *Kcnn3* | -0.21 | 2.07E-06 | *Bckdk* | -1.01 | 4.1E-04 |
| *Snrk* | -0.35 | 1.1E-02 | -0.49 | 1.1E-06 | *Tmed5* | -0.21 | 8.35E-04 | *C77080* | -0.99 | 5.9E-05 |
| *Zfp608* | -0.34 | 2.4E-12 | -0.43 | 3.8E-17 | *Trps1* | -0.20 | 1.99E-04 | *Gm2000* | -0.98 | 6.1E-05 |
| *Tsc22d1* | -0.33 | 1.4E-03 | -0.59 | 2.4E-11 | *Ywhah* | -0.18 | 1.26E-03 | *Cd36* | -0.95 | 4.6E-25 |
| *Peli1* | -0.33 | 1.2E-05 | -0.39 | 1.9E-08 | *Cytip* | -0.17 | 5.67E-03 | *Ctdsp2* | -0.91 | 3.6E-07 |
| *Prkar2b* | -0.32 | 1.1E-11 | -0.40 | 4.6E-15 | *Ikzf1* | -0.17 | 4.90E-02 | *Sdc3* | -0.90 | 6.1E-17 |
| *Sik3* | -0.31 | 2.9E-02 | -0.51 | 2.8E-07 | *Sat1* | -0.16 | 2.54E-04 | *Trim28* | -0.88 | 1.4E-02 |
| *Tmem154* | -0.31 | 9.0E-03 | -0.35 | 1.3E-02 | *Ahnak* | -0.14 | 3.24E-02 | *Rbpms* | -0.88 | 2.5E-03 |
| *Ear1* | -0.30 | 9.4E-05 | -0.51 | 2.5E-16 | *Lyn* | -0.14 | 4.64E-05 | *Acot1* | -0.87 | 3.6E-07 |

\*A complete list of differentially expressed genes is included in Supplemental Table S8.

AM cluster 2, which accounts for ∼16% (937 out of 5870; **Supplemental Table S??**) of total annotated AMs, was analyzed to identify DEGs between 1 ppm O_3_ versus FA and 1.5 ppm O_3_ versus FA group. Using the cut-off criteria (|average log2FC| ≥ 0.1, adjusted *P* value ≤ 0.05, min.pct ≥ 0.1), the analyses identified 225 DEGs (upregulated, 159; downregulated, 66) in 1 ppm O_3_ versus FA, and 599 DEGs (upregulated, 301; downregulated, 298) in 1.5 ppm O_3_ versus FA groups (**Figure 6E, Supplemental Table S9**). Among the DEGs identified by differential expression analyses, a total of 98 DEGs (upregulated, 67; downregulated, 31) were common to 1 ppm and 1.5 ppm O_3_ with respect to FA (**Figure 6F**, **Table 4, Supplemental Table S9**). The IPA revealed that these 98 DEGs were associated with the enrichment of pathways involved in protein translation (**Figure 7C, Supplemental Table S12**). We identified 124 DEGs (upregulated, 91; downregulated, 33), and 498 DEGs (upregulated, 232; downregulated, 266) that were uniquely expressed in 1 ppm O_3_ versus FA and 1.5 ppm O_3_ versus FA, respectively (**Figure 6F**, **Table 4, Supplemental Table S9**). While the unique DEGs in 1 ppm O_3_ versus FA comparison were involved in the enrichment of protein translation pathways, the unique DEGs in 1.5 ppm O_3_ versus FA comparisons were involved in the enrichment of pathways involved in mitochondrial function as well as protein translation (**Figure 7C, Supplemental Table S12**). The unique DEGs in 1.5 ppm O_3_ versus FA comparison were involved in the suppression of mitochondrial dysfunction pathways (**Figure 7C, Supplemental Table S12**).

**Table 4.**
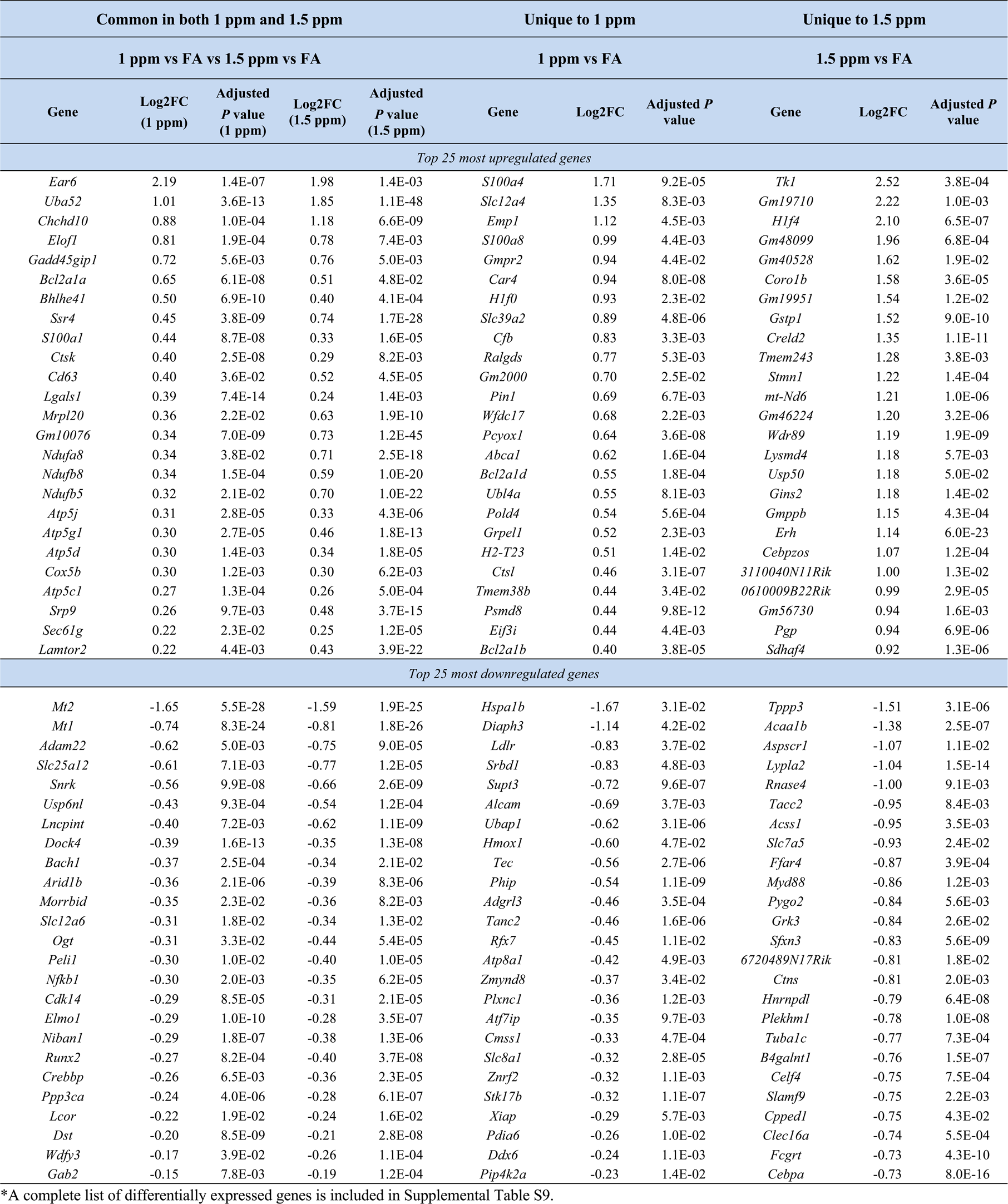
Top 25 most upregulated and top 25 most downregulated genes in Cluster 2 AMs from 1 ppm O_3_ and 1.5 ppm O_3_ (common and unique) with respect to filtered air (FA) 2.

| Common in both 1 ppm and 1.5 ppm |  |  |  |  | Unique to 1 ppm |  |  | Unique to 1.5 ppm |  |  |
| --- | --- | --- | --- | --- | --- | --- | --- | --- | --- | --- |
| 1 ppm vs FA vs 1.5 ppm vs FA |  |  |  |  | 1 ppm vs FA |  |  | 1.5 ppm vs FA |  |  |
| Gene | Log2FC<br>(1 ppm) | Adjusted<br>P value<br>(1 ppm) | Log2FC<br>(1.5 ppm) | Adjusted<br>P value<br>(1.5 ppm) | Gene | Log2FC | Adjusted P<br>value | Gene | Log2FC | Adjusted P<br>value |
| Top 25 most upregulated genes |  |  |  |  |  |  |  |  |  |  |
| *Ear6* | 2.19 | 1.4E-07 | 1.98 | 1.4E-03 | *Sl00a4* | 1.71 | 9.2E-05 | *Tkl* | 2.52 | 3.8E-04 |
| *Uba52* | 1.01 | 3.6E-13 | 1.85 | 1.1E-48 | *Slc12a4* | 1.35 | 8.3E-03 | *Gm19710* | 2.22 | 1.0E-03 |
| *Chchd10* | 0.88 | 1.0E-04 | 1.18 | 6.6E-09 | *Emp1* | 1.12 | 4.5E-03 | *Hlf4* | 2.10 | 6.5E-07 |
| *Elofl* | 0.81 | 1.9E-04 | 0.78 | 7.4E-03 | *Sl00a8* | 0.99 | 4.4E-03 | *Gm48099* | 1.96 | 6.8E-04 |
| *Gadd45gip1* | 0.72 | 5.6E-03 | 0.76 | 5.0E-03 | *Gmpr2* | 0.94 | 4.4E-02 | *Gm40528* | 1.62 | 1.9E-02 |
| *Bcl2a1a* | 0.65 | 6.1E-08 | 0.51 | 4.8E-02 | *Car4* | 0.94 | 8.0E-08 | *Coro1b* | 1.58 | 3.6E-05 |
| *Bhlhe41* | 0.50 | 6.9E-10 | 0.40 | 4.1E-04 | *Hlf0* | 0.93 | 2.3E-02 | *Gm19951* | 1.54 | 1.2E-02 |
| *Ssr4* | 0.45 | 3.8E-09 | 0.74 | 1.7E-28 | *Slc39a2* | 0.89 | 4.8E-06 | *Gstp1* | 1.52 | 9.0E-10 |
| *Sl00a1* | 0.44 | 8.7E-08 | 0.33 | 1.6E-05 | *Cfb* | 0.83 | 3.3E-03 | *Creld2* | 1.35 | 1.1E-11 |
| *Ctsk* | 0.40 | 2.5E-08 | 0.29 | 8.2E-03 | *Ralgds* | 0.77 | 5.3E-03 | *Tmem243* | 1.28 | 3.8E-03 |
| *Cd63* | 0.40 | 3.6E-02 | 0.52 | 4.5E-05 | *Gm2000* | 0.70 | 2.5E-02 | *Stmn1* | 1.22 | 1.4E-04 |
| *Lgals1* | 0.39 | 7.4E-14 | 0.24 | 1.4E-03 | *Pin1* | 0.69 | 6.7E-03 | *mt-Nd6* | 1.21 | 1.0E-06 |
| *Mrpl20* | 0.36 | 2.2E-02 | 0.63 | 1.9E-10 | *Wfdc17* | 0.68 | 2.2E-03 | *Gm46224* | 1.20 | 3.2E-06 |
| *Gm10076* | 0.34 | 7.0E-09 | 0.73 | 1.2E-45 | *Pcyox1* | 0.64 | 3.6E-08 | *Wdr89* | 1.19 | 1.9E-09 |
| *Ndufa8* | 0.34 | 3.8E-02 | 0.71 | 2.5E-18 | *Abca1* | 0.62 | 1.6E-04 | *Lysmd4* | 1.18 | 5.7E-03 |
| *Ndufb8* | 0.34 | 1.5E-04 | 0.59 | 1.0E-20 | *Bcl2a1d* | 0.55 | 1.8E-04 | *Usp50* | 1.18 | 5.0E-02 |
| *Ndufb5* | 0.32 | 2.1E-02 | 0.70 | 1.0E-22 | *Ubl4a* | 0.55 | 8.1E-03 | *Gins2* | 1.18 | 1.4E-02 |
| *Atp5j* | 0.31 | 2.8E-05 | 0.33 | 4.3E-06 | *Pold4* | 0.54 | 5.6E-04 | *Gmppb* | 1.15 | 4.3E-04 |
| *Atp5g1* | 0.30 | 2.7E-05 | 0.46 | 1.8E-13 | *Grpel1* | 0.52 | 2.3E-03 | *Erh* | 1.14 | 6.0E-23 |
| *Atp5d* | 0.30 | 1.4E-03 | 0.34 | 1.8E-05 | *H2-T23* | 0.51 | 1.4E-02 | *Cebpzos* | 1.07 | 1.2E-04 |
| *Cox5b* | 0.30 | 1.2E-03 | 0.30 | 6.2E-03 | *Ctsl* | 0.46 | 3.1E-07 | *3110040N11Rik* | 1.00 | 1.3E-02 |
| *Atp5c1* | 0.27 | 1.3E-04 | 0.26 | 5.0E-04 | *Tmem38b* | 0.44 | 3.4E-02 | *0610009B22Rik* | 0.99 | 2.9E-05 |
| *Srp9* | 0.26 | 9.7E-03 | 0.48 | 3.7E-15 | *Psmc8* | 0.44 | 9.8E-12 | *Gm56730* | 0.94 | 1.6E-03 |
| *Sec61g* | 0.22 | 2.3E-02 | 0.25 | 1.2E-05 | *Eif3i* | 0.44 | 4.4E-03 | *Pgp* | 0.94 | 6.9E-06 |
| *Lamtor2* | 0.22 | 4.4E-03 | 0.43 | 3.9E-22 | *Bcl2a1b* | 0.40 | 3.8E-05 | *Sdhaf4* | 0.92 | 1.3E-06 |
| Top 25 most downregulated genes |  |  |  |  |  |  |  |  |  |  |
| *Mt2* | -1.65 | 5.5E-28 | -1.59 | 1.9E-25 | *Hspa1b* | -1.67 | 3.1E-02 | *Tppp3* | -1.51 | 3.1E-06 |
| *Mt1* | -0.74 | 8.3E-24 | -0.81 | 1.8E-26 | *Diaph3* | -1.14 | 4.2E-02 | *Acaa1b* | -1.38 | 2.5E-07 |
| *Adam22* | -0.62 | 5.0E-03 | -0.75 | 9.0E-05 | *Ldlr* | -0.83 | 3.7E-02 | *Aspscr1* | -1.07 | 1.1E-02 |
| *Slc25a12* | -0.61 | 7.1E-03 | -0.77 | 1.2E-05 | *Srbdl1* | -0.83 | 4.8E-03 | *Lypla2* | -1.04 | 1.5E-14 |
| *Snrk* | -0.56 | 9.9E-08 | -0.66 | 2.6E-09 | *Supt3* | -0.72 | 9.6E-07 | *Rnase4* | -1.00 | 9.1E-03 |
| *Usp6nl* | -0.43 | 9.3E-04 | -0.54 | 1.2E-04 | *Alcam* | -0.69 | 3.7E-03 | *Tace2* | -0.95 | 8.4E-03 |
| *Lncpint* | -0.40 | 7.2E-03 | -0.62 | 1.1E-09 | *Ubap1* | -0.62 | 3.1E-06 | *Acss1* | -0.95 | 3.5E-03 |
| *Dock4* | -0.39 | 1.6E-13 | -0.35 | 1.3E-08 | *Hmox1* | -0.60 | 4.7E-02 | *Slc7a5* | -0.93 | 2.4E-02 |
| *Bach1* | -0.37 | 2.5E-04 | -0.34 | 2.1E-02 | *Tec* | -0.56 | 2.7E-06 | *Ffar4* | -0.87 | 3.9E-04 |
| *Arid1b* | -0.36 | 2.1E-06 | -0.39 | 8.3E-06 | *Phip* | -0.54 | 1.1E-09 | *Myd88* | -0.86 | 1.2E-03 |
| *Morrbid* | -0.35 | 2.3E-02 | -0.36 | 8.2E-03 | *Adgrl3* | -0.46 | 3.5E-04 | *Pygo2* | -0.84 | 5.6E-03 |
| *Slc12a6* | -0.31 | 1.8E-02 | -0.34 | 1.3E-02 | *Tanc2* | -0.46 | 1.6E-06 | *Grk3* | -0.84 | 2.6E-02 |
| *Ogt* | -0.31 | 3.3E-02 | -0.44 | 5.4E-05 | *Rfx7* | -0.45 | 1.1E-02 | *Sfxn3* | -0.83 | 5.6E-09 |
| *Peli1* | -0.30 | 1.0E-02 | -0.40 | 1.0E-05 | *Atp8a1* | -0.42 | 4.9E-03 | *6720489N17Rik* | -0.81 | 1.8E-02 |
| *Nfkb1* | -0.30 | 2.0E-03 | -0.35 | 6.2E-05 | *Zmynd8* | -0.37 | 3.4E-02 | *Ctns* | -0.81 | 2.0E-03 |
| *Cdk14* | -0.29 | 8.5E-05 | -0.31 | 2.1E-05 | *Plxnc1* | -0.36 | 1.2E-03 | *Hnrnpdl* | -0.79 | 6.4E-08 |
| *Elmo1* | -0.29 | 1.0E-10 | -0.28 | 3.5E-07 | *Atf7ip* | -0.35 | 9.7E-03 | *Plekhm1* | -0.78 | 1.0E-08 |
| *Niban1* | -0.29 | 1.8E-07 | -0.38 | 1.3E-06 | *Cmss1* | -0.33 | 4.7E-04 | *Tuba1c* | -0.77 | 7.3E-04 |
| *Runx2* | -0.27 | 8.2E-04 | -0.40 | 3.7E-08 | *Slc8a1* | -0.32 | 2.8E-05 | *B4galnt1* | -0.76 | 1.5E-07 |
| *Crebbp* | -0.26 | 6.5E-03 | -0.36 | 2.3E-05 | *Znrf2* | -0.32 | 1.1E-03 | *Celf4* | -0.75 | 7.5E-04 |
| *Ppp3ca* | -0.24 | 4.0E-06 | -0.28 | 6.1E-07 | *Stk17b* | -0.32 | 1.1E-07 | *Slamf9* | -0.75 | 2.2E-03 |
| *Lcor* | -0.22 | 1.9E-02 | -0.24 | 1.6E-02 | *Xiap* | -0.29 | 5.7E-03 | *Cpped1* | -0.75 | 4.3E-02 |
| *Dst* | -0.20 | 8.5E-09 | -0.21 | 2.8E-08 | *Pdia6* | -0.26 | 1.0E-02 | *Clec16a* | -0.74 | 5.5E-04 |
| *Wdfy3* | -0.17 | 3.9E-02 | -0.26 | 1.1E-04 | *Ddx6* | -0.24 | 1.1E-03 | *Fcgrt* | -0.73 | 4.3E-10 |
| *Gab2* | -0.15 | 7.8E-03 | -0.19 | 1.2E-04 | *Pip4k2a* | -0.23 | 1.4E-02 | *Cebpa* | -0.73 | 8.0E-16 |

\*A complete list of differentially expressed genes is included in Supplemental Table S9.

AM cluster 3, which accounts for ∼15% (879 out of 5870; **Supplemental Table S5**) of total annotated AMs, was analyzed to identify DEGs between 1 ppm O_3_ versus FA and 1.5 ppm O_3_ versus FA group. Using the cut-off criteria (|average log2FC| ≥ 0.1, adjusted *P* value ≤ 0.05, min.pct ≥ 0.1), the analyses revealed 369 DEGs (upregulated, 206 downregulated, 163) in 1 ppm O_3_ versus FA, and 834 DEGs (upregulated, 447; downregulated, 387) in 1.5 ppm O_3_ versus FA (**Figure 6G, Supplemental Table S10**). Among the DEGs identified by differential expression analyses, a total of 217 DEGs (upregulated, 125; downregulated, 92) were common to 1 ppm and 1.5 ppm O_3_ with respect to FA (**Figure 6H**, **Table 5, Supplemental Table S10**). These 217 DEGs were associated with the enrichment of pathways involved in protein translation and proteasome-mediated degradation pathways involving genes such as *Psma3, Psmb1, Psmb2, Psmb8, Rbx1m Rps27a, Sem1*, *Uba52* (**Figure 7D, Supplemental Table S10 and S12**). On the other hand, we identified 152 DEGs (upregulated, 81; downregulated, 71), and 617 DEGs (upregulated, 322; downregulated, 295) that were uniquely expressed in 1 ppm O_3_ versus FA and 1.5 ppm O_3_ versus FA, respectively (**Figure 6H**, **Table 5, Supplemental Table S10**).While the unique DEGs in 1 ppm O_3_ versus FA comparison were involved in the enrichment of protein translation pathways, the unique DEGs in 1.5 ppm O_3_ versus FA comparison were involved in the enrichment of pathways, including cell-cycle, mitochondrial biogenesis, and ubiquitination (**Figure 7D, Supplemental Table S12**). The unique DEGs in 1.5 ppm O_3_ versus FA comparison were involved in the suppression of mitochondrial dysfunction and macrophage alternative activation signaling pathway (**Figure 7D, Supplemental Table S12**).

**Table 5.** Top 25 most upregulated and top 25 most downregulated genes in Cluster 3 AMs from 1 ppm O_3_ and 1.5 ppm O_3_ (common and unique) with respect to filtered air (FA).

| Common in both 1 ppm and 1.5 ppm |  |  |  |  | Unique to 1 ppm |  |  | Unique to 1.5 ppm |  |  |
| --- | --- | --- | --- | --- | --- | --- | --- | --- | --- | --- |
| 1 ppm vs FA vs 1.5 ppm vs FA |  |  |  |  | 1 ppm vs FA |  |  | 1.5 ppm vs FA |  |  |
| Gene | Log2FC<br>(1 ppm) | Adjusted<br>P value<br>(1 ppm) | Log2FC<br>(1.5 ppm) | Adjusted<br>P value<br>(1.5 ppm) | Gene | Log2FC | Adjusted P<br>value | Gene | Log2FC | Adjusted P<br>value |
| Top 25 most upregulated genes |  |  |  |  |  |  |  |  |  |  |
| *Wfdc17* | 1.02 | 1.2E-04 | 0.78 | 3.4E-03 | *Cjfb* | 1.25 | 5.6E-03 | *5430427M07Rik* | 3.27 | 9.0E-04 |
| *Uba52* | 0.96 | 6.7E-07 | 2.00 | 2.1E-55 | *Dtd2* | 1.09 | 9.7E-03 | *H3c7* | 2.64 | 3.3E-02 |
| *Car4* | 0.85 | 1.6E-08 | 0.67 | 3.4E-06 | *Scd2* | 0.99 | 1.4E-05 | *H2ac10* | 2.19 | 4.1E-04 |
| *Bbln* | 0.74 | 9.0E-05 | 0.74 | 3.7E-05 | *Emilin2* | 0.83 | 8.0E-03 | *H4c8* | 2.16 | 1.6E-08 |
| *Fn1* | 0.63 | 9.9E-06 | 0.59 | 8.1E-07 | *Pin1* | 0.77 | 3.5E-02 | *H2ac18* | 2.16 | 6.5E-06 |
| *Mrpl52* | 0.62 | 7.2E-05 | 0.48 | 2.4E-02 | *Clec12a* | 0.76 | 2.6E-02 | *Hspa1b* | 2.03 | 4.4E-10 |
| *Mif* | 0.59 | 3.5E-04 | 0.59 | 4.6E-06 | *Slirp* | 0.71 | 3.0E-02 | *H3c6* | 2.02 | 2.2E-06 |
| *Ufc1* | 0.59 | 6.0E-03 | 0.69 | 3.9E-09 | *Eif4ebp1* | 0.70 | 1.5E-02 | *H3c2* | 1.98 | 2.5E-02 |
| *Dnajc19* | 0.58 | 1.4E-03 | 0.76 | 2.6E-13 | *Hypk* | 0.67 | 7.5E-05 | *Gstp1* | 1.88 | 1.9E-15 |
| *Park7* | 0.54 | 8.2E-03 | 0.71 | 4.4E-11 | *Psmd6* | 0.61 | 2.9E-02 | *H2bc11* | 1.88 | 1.8E-02 |
| *Ndufb6* | 0.53 | 2.2E-03 | 0.50 | 2.8E-05 | *Mrpl30* | 0.56 | 4.8E-03 | *H3c3* | 1.69 | 1.0E-05 |
| *Uqcrq* | 0.46 | 4.7E-12 | 0.30 | 3.9E-06 | *Atp5md* | 0.56 | 2.0E-08 | *Gm40528* | 1.64 | 3.0E-05 |
| *Chmp2a* | 0.44 | 2.6E-03 | 0.43 | 4.3E-05 | *Wfdc21* | 0.52 | 4.5E-07 | *H1f4* | 1.59 | 3.4E-16 |
| *Lamtor4* | 0.44 | 4.7E-04 | 0.41 | 2.8E-05 | *Ctsl* | 0.51 | 1.8E-04 | *H4c4* | 1.57 | 4.4E-10 |
| *Ndufb8* | 0.44 | 5.3E-08 | 0.52 | 5.8E-20 | *Ndufa13* | 0.48 | 1.1E-04 | *H2ac24* | 1.55 | 8.0E-15 |
| *Psmd8* | 0.44 | 8.4E-06 | 0.41 | 1.0E-06 | *Cd52* | 0.43 | 1.7E-04 | *H2ac4* | 1.53 | 2.6E-04 |
| *Atp5g1* | 0.44 | 1.9E-09 | 0.48 | 4.6E-17 | *Rac2* | 0.42 | 4.9E-02 | *Gm48099* | 1.50 | 1.2E-02 |
| *Ndufb9* | 0.43 | 6.8E-07 | 0.56 | 1.9E-21 | *Micos13* | 0.41 | 4.9E-02 | *H4c3* | 1.49 | 1.5E-02 |
| *Ssr4* | 0.43 | 1.6E-04 | 0.74 | 1.3E-28 | *Gngt2* | 0.41 | 1.8E-05 | *H2ac8* | 1.47 | 5.3E-10 |
| *Ndufc1* | 0.43 | 1.3E-02 | 0.42 | 1.9E-04 | *Uqcrb* | 0.41 | 6.0E-08 | *H2ac20* | 1.46 | 4.1E-06 |
| *Cenpx* | 0.42 | 3.3E-02 | 0.54 | 4.0E-10 | *Atp5mpl* | 0.40 | 9.3E-08 | *Usp50* | 1.44 | 5.3E-04 |
| *Swi5* | 0.42 | 4.3E-03 | 0.49 | 9.2E-10 | *Elob* | 0.39 | 6.4E-08 | *Ube2s* | 1.41 | 4.5E-06 |
| *Timm10b* | 0.42 | 1.3E-04 | 0.47 | 1.4E-11 | *Tmem258* | 0.39 | 1.5E-02 | *Ccr5* | 1.35 | 2.4E-02 |
| *Spcs2* | 0.42 | 9.0E-04 | 0.31 | 3.2E-02 | *Tmem256* | 0.39 | 4.5E-07 | *H1f5* | 1.29 | 2.5E-14 |
| *Gm10076* | 0.41 | 7.8E-12 | 0.70 | 6.7E-44 | *Uqcr11* | 0.38 | 1.2E-06 | *Chchd10* | 1.28 | 1.7E-06 |
| Top 25 most downregulated genes |  |  |  |  |  |  |  |  |  |  |
| *Hbb-bs* | -3.86 | 1.5E-03 | -2.59 | 1.2E-03 | *Mybl1* | -1.05 | 3.9E-02 | *Rbfox1* | -1.81 | 8.8E-05 |
| *Dusp6* | -0.98 | 1.4E-03 | -0.92 | 1.3E-04 | *Reps2* | -0.95 | 1.8E-08 | *Acaa1b* | -1.59 | 8.6E-18 |
| *Rnd3* | -0.72 | 2.3E-02 | -0.71 | 2.5E-03 | *Mcm9* | -0.77 | 2.6E-02 | *Gm49980* | -1.57 | 8.9E-03 |
| *N4bp2l1* | -0.71 | 1.2E-05 | -0.46 | 2.8E-02 | *Sclt1* | -0.70 | 8.6E-03 | *Gm32849* | -1.57 | 1.2E-03 |
| *Slc9a7* | -0.66 | 9.8E-08 | -0.65 | 1.3E-10 | *Plxnc1* | -0.67 | 7.4E-09 | *Slc7a5* | -1.32 | 2.3E-07 |
| *Tll5* | -0.61 | 1.4E-02 | -0.54 | 7.3E-03 | *Brip1* | -0.65 | 1.7E-02 | *Mt2* | -1.26 | 8.0E-05 |
| *Adam22* | -0.60 | 5.3E-03 | -0.93 | 4.5E-12 | *Mast2* | -0.61 | 1.1E-03 | *Gdi1* | -1.21 | 1.0E-03 |
| *Zfp609* | -0.57 | 2.1E-02 | -0.53 | 3.9E-03 | *Tec* | -0.60 | 8.9E-04 | *Pltp* | -1.07 | 2.0E-05 |
| *Patj* | -0.55 | 1.1E-05 | -0.91 | 3.1E-21 | *Gm47283* | -0.60 | 2.0E-03 | *Rnase4* | -1.05 | 6.4E-04 |
| *Vkorc1l1* | -0.51 | 6.8E-08 | -0.36 | 7.5E-05 | *Supt3* | -0.56 | 3.1E-02 | *Klhl21* | -1.03 | 7.1E-03 |
| *Dym* | -0.51 | 1.5E-03 | -0.50 | 4.8E-05 | *Kdm3b* | -0.54 | 7.6E-04 | *B4galnt1* | -1.03 | 4.1E-25 |
| *Ccpgl* | -0.51 | 9.0E-05 | -0.62 | 3.3E-09 | *Tanc2* | -0.53 | 7.0E-06 | *Themis2* | -1.00 | 4.5E-11 |
| *Banp* | -0.50 | 4.0E-02 | -0.53 | 4.3E-05 | *Rapgef2* | -0.53 | 9.3E-03 | *Cd83* | -0.94 | 1.0E-02 |
| *Lncpint* | -0.50 | 7.8E-05 | -0.67 | 2.2E-13 | *Ezh2* | -0.51 | 1.1E-04 | *Eeig2* | -0.92 | 3.7E-04 |
| *Bckdhb* | -0.49 | 8.2E-03 | -0.62 | 9.0E-07 | *Rsbn1l* | -0.51 | 2.9E-02 | *Marveld2* | -0.92 | 3.8E-04 |
| *Lpin1* | -0.48 | 3.8E-06 | -0.61 | 2.6E-14 | *Rb1* | -0.50 | 1.5E-06 | *Naa30* | -0.91 | 4.2E-02 |
| *Cpne5* | -0.47 | 6.8E-04 | -0.50 | 2.2E-06 | *Bach1* | -0.50 | 5.4E-04 | *Tafazzin* | -0.90 | 3.1E-04 |
| *Slc10a7* | -0.43 | 1.3E-02 | -0.38 | 2.4E-02 | *Nsd2* | -0.48 | 3.2E-04 | *Acss1* | -0.89 | 1.8E-08 |
| *Tsc22d1* | -0.43 | 2.8E-02 | -0.52 | 9.0E-06 | *Pbx3* | -0.47 | 1.9E-02 | *Lypla2* | -0.88 | 3.3E-11 |
| *Peli1* | -0.43 | 2.1E-07 | -0.44 | 2.5E-11 | *Pola1* | -0.46 | 3.2E-03 | *Cebpa* | -0.85 | 5.1E-16 |
| *Ccn1l* | -0.43 | 3.5E-03 | -0.43 | 2.1E-05 | *Marf1* | -0.46 | 2.8E-02 | *Phlda1* | -0.85 | 1.4E-04 |
| *Lrp12* | -0.41 | 3.0E-06 | -0.39 | 7.6E-08 | *Bmp2k* | -0.46 | 1.3E-02 | *Fcgrt* | -0.84 | 1.1E-16 |
| *Snrk* | -0.41 | 2.7E-03 | -0.55 | 3.3E-09 | *Kmt2c* | -0.43 | 3.8E-05 | *Hsd17b11* | -0.84 | 4.1E-05 |
| *Abhd17c* | -0.40 | 3.0E-02 | -0.40 | 1.1E-03 | *Zbtb20* | -0.42 | 5.5E-04 | *Dcun1d3* | -0.83 | 1.5E-05 |
| *Kcnn3* | -0.39 | 2.2E-07 | -0.24 | 9.6E-04 | *Xpo7* | -0.41 | 1.9E-02 | *Cdh1* | -0.83 | 1.6E-03 |

\*A complete list of differentially expressed genes is included in Supplemental Table S10.

AM cluster 4, which accounts for ∼7% (422 out of 5870; **Supplemental Table S5**) of total annotated AMs, was analyzed to identify DEGs between 1 ppm O_3_ versus FA and 1.5 ppm O_3_ versus FA group. Using the cut-off criteria (|average log2FC| ≥ 0.1, adjusted *P* value ≤ 0.05, min.pct ≥ 0.1), the analyses identified 39 DEGs (upregulated, 28; downregulated, 11) in 1 ppm O_3_ versus FA, and 28 DEGs (upregulated, 21; downregulated, 7) in 1.5 ppm O_3_ versus FA (**Figure 6I, Supplemental Table S11)**. Among the DEGs identified by differential expression analyses, a total of 4 DEGs (upregulated, 1; downregulated, 3) were common to 1 ppm and 1.5 ppm O_3_ with respect to FA (**Figure 6J**, **Table 6, Supplemental Table S11**). On the other hand, we identified 35 DEGs (upregulated, 27; downregulated, 8), and 24 DEGs (upregulated, 6; downregulated, 18) that were uniquely expressed in 1 ppm O_3_ versus FA and 1.5 ppm O_3_ versus FA, respectively (**Figure 6J**, **Table 6, Supplemental Table S11**). The IP analyses revealed that these unique and common DEGs were not influencing biological pathways indicating an AM subpopulation that is minimally affected by O_3_ exposure.

**Table 6.** Top 25 most upregulated and top 25 most downregulated genes in Cluster 4 AMs from 1 ppm O_3_ and 1.5 ppm O_3_ (common and unique) with respect to filtered air (FA).

| Common in both 1 ppm and 1.5 ppm |  |  |  |  | Unique to 1 ppm |  |  | Unique to 1.5 ppm |  |  |
| --- | --- | --- | --- | --- | --- | --- | --- | --- | --- | --- |
| 1 ppm vs FA vs 1.5 ppm vs FA |  |  |  |  | 1 ppm vs FA |  |  | 1.5 ppm vs FA |  |  |
| Gene | Log2FC<br>(1 ppm) | Adjusted<br>P value<br>(1 ppm) | Log2FC<br>(1.5 ppm) | Adjusted<br>P value<br>(1.5 ppm) | Gene | Log2FC | Adjusted P<br>value | Gene | Log2FC | Adjusted P<br>value |
| Top 25 most upregulated genes |  |  |  |  |  |  |  |  |  |  |
| *Fth1* | 0.26 | 9.4E-05 | 0.27 | 5.1E-07 | *Rpl29* | 1.05 | 9.2E-03 | *mt-Nd6* | 0.80 | 4.8E-02 |
|  |  |  |  |  | *Rps27* | 0.98 | 7.0E-03 | *mt-Atp8* | 0.78 | 1.8E-02 |
|  |  |  |  |  | *Rpl28* | 0.82 | 2.0E-02 | *mt-Nd4l* | 0.57 | 3.6E-16 |
|  |  |  |  |  | *Rpl18a* | 0.78 | 2.4E-03 | *Ppia* | 0.56 | 4.1E-02 |
|  |  |  |  |  | *Rps11* | 0.77 | 3.9E-04 | *Lyst* | 0.47 | 1.6E-02 |
|  |  |  |  |  | *Rpl10a* | 0.76 | 3.5E-02 | *mt-Nd5* | 0.34 | 1.6E-07 |
|  |  |  |  |  | *Rpl19* | 0.74 | 3.3E-03 |  |  |  |
|  |  |  |  |  | *Rps5* | 0.74 | 3.5E-02 |  |  |  |
|  |  |  |  |  | *Rpl6* | 0.73 | 2.7E-02 |  |  |  |
|  |  |  |  |  | *Rps13* | 0.72 | 1.7E-02 |  |  |  |
|  |  |  |  |  | *Rpl30* | 0.70 | 3.6E-04 |  |  |  |
|  |  |  |  |  | *Rpl13a* | 0.67 | 2.1E-02 |  |  |  |
|  |  |  |  |  | *Rps23* | 0.66 | 4.8E-03 |  |  |  |
|  |  |  |  |  | *Serinc3* | 0.66 | 5.3E-03 |  |  |  |
|  |  |  |  |  | *Rpl10* | 0.65 | 3.2E-02 |  |  |  |
|  |  |  |  |  | *Rps10* | 0.61 | 3.6E-03 |  |  |  |
|  |  |  |  |  | *Rps16* | 0.57 | 9.0E-03 |  |  |  |
|  |  |  |  |  | *Fcer1g* | 0.55 | 2.0E-05 |  |  |  |
|  |  |  |  |  | *Rpl13* | 0.55 | 2.4E-03 |  |  |  |
|  |  |  |  |  | *Fkbp5* | 0.53 | 1.5E-03 |  |  |  |
|  |  |  |  |  | *Rps8* | 0.48 | 1.6E-02 |  |  |  |
|  |  |  |  |  | *Rpl23* | 0.46 | 4.0E-03 |  |  |  |
|  |  |  |  |  | *Itm2b* | 0.34 | 1.2E-02 |  |  |  |
|  |  |  |  |  | *Lrp1* | 0.30 | 2.2E-03 |  |  |  |
|  |  |  |  |  | *Ctss* | 0.21 | 3.7E-04 |  |  |  |
| Top 25 most downregulated genes |  |  |  |  |  |  |  |  |  |  |
| *Mt2* | -3.20 | 2.0E-05 | -1.83 | 2.7E-03 | *Tbc1d2b* | -2.33 | 3.1E-02 | *Gm32849* | -1.72 | 8.0E-04 |
| *Bckdhb* | -0.50 | 1.5E-02 | -0.57 | 3.3E-02 | *Pparg* | -0.34 | 3.2E-04 | *Acaa1b* | -1.55 | 2.5E-03 |
| *Gab2* | -0.20 | 4.9E-07 | -0.15 | 3.2E-04 | *Zfp608* | -0.33 | 2.0E-02 | *Scgb1a1* | -1.37 | 4.8E-02 |
|  |  |  |  |  | *Ankrd44* | -0.26 | 3.8E-03 | *Cd36* | -0.92 | 9.4E-03 |
|  |  |  |  |  | *Kcnn3* | -0.26 | 5.3E-04 | *Jarid2* | -0.84 | 5.6E-04 |
|  |  |  |  |  | *Dock4* | -0.20 | 4.4E-03 | *Lepr* | -0.79 | 1.6E-02 |
|  |  |  |  |  | *mt-Nd4* | -0.17 | 5.6E-05 | *Rin2* | -0.77 | 2.5E-02 |
|  |  |  |  |  | *Dock10* | -0.10 | 3.0E-03 | *Adam22* | -0.61 | 3.2E-02 |
|  |  |  |  |  |  |  |  | *Patj* | -0.61 | 5.6E-03 |
|  |  |  |  |  |  |  |  | *Ccnd3* | -0.54 | 4.4E-04 |
|  |  |  |  |  |  |  |  | *Tmbim6* | -0.48 | 1.4E-02 |
|  |  |  |  |  |  |  |  | *Lima1* | -0.46 | 3.8E-02 |
|  |  |  |  |  |  |  |  | *Laptm5* | -0.45 | 5.4E-03 |
|  |  |  |  |  |  |  |  | *mt-Nd3* | -0.36 | 1.5E-15 |
|  |  |  |  |  |  |  |  | *Foxn3* | -0.29 | 8.7E-04 |
|  |  |  |  |  |  |  |  | *Cracd* | -0.29 | 6.8E-05 |
|  |  |  |  |  |  |  |  | *Nav2* | -0.24 | 2.0E-02 |
|  |  |  |  |  |  |  |  | *mt-Cytb* | -0.16 | 5.4E-13 |

\*A complete list of differentially expressed genes is included in Supplemental Table S11.

### Ozone exposure modulates expression of genes relevant to macrophage activation pathways

To assess the activation status of AMs from different treatment groups, first, we randomly selected 50 AMs from FA-exposed, 1 ppm O_3_-exposed, and 1.5 ppm O_3_-exposed mice and compared the expression of genes relevant to M1 and M2 activation responses. Compared to the AMs from FA-exposed mice, AMs from mice exposed to 1 ppm O_3_ displayed increased expression of genes associated with both M1 and M2 activation responses (**Figure 8A**). In contrast, AMs from mice exposed to 1.5 ppm O_3_ exhibited decreased expression of genes related to both M1 and M2 activation responses (**Figure 8A**). However, the comparisons of M1 and M2 activation signatures at a single-cell level revealed increased expression of genes relevant to both M1 and M2 activation. For instance, cells corresponding to asterisks in **Figure 8A** suggested higher copy numbers of transcripts relevant to M1, e.g., *Il18*, *Cd68, Aldoa,* as well as M2, e.g., *Chil3, Clec7a, Trem2,* activation.

**Figure 8:**
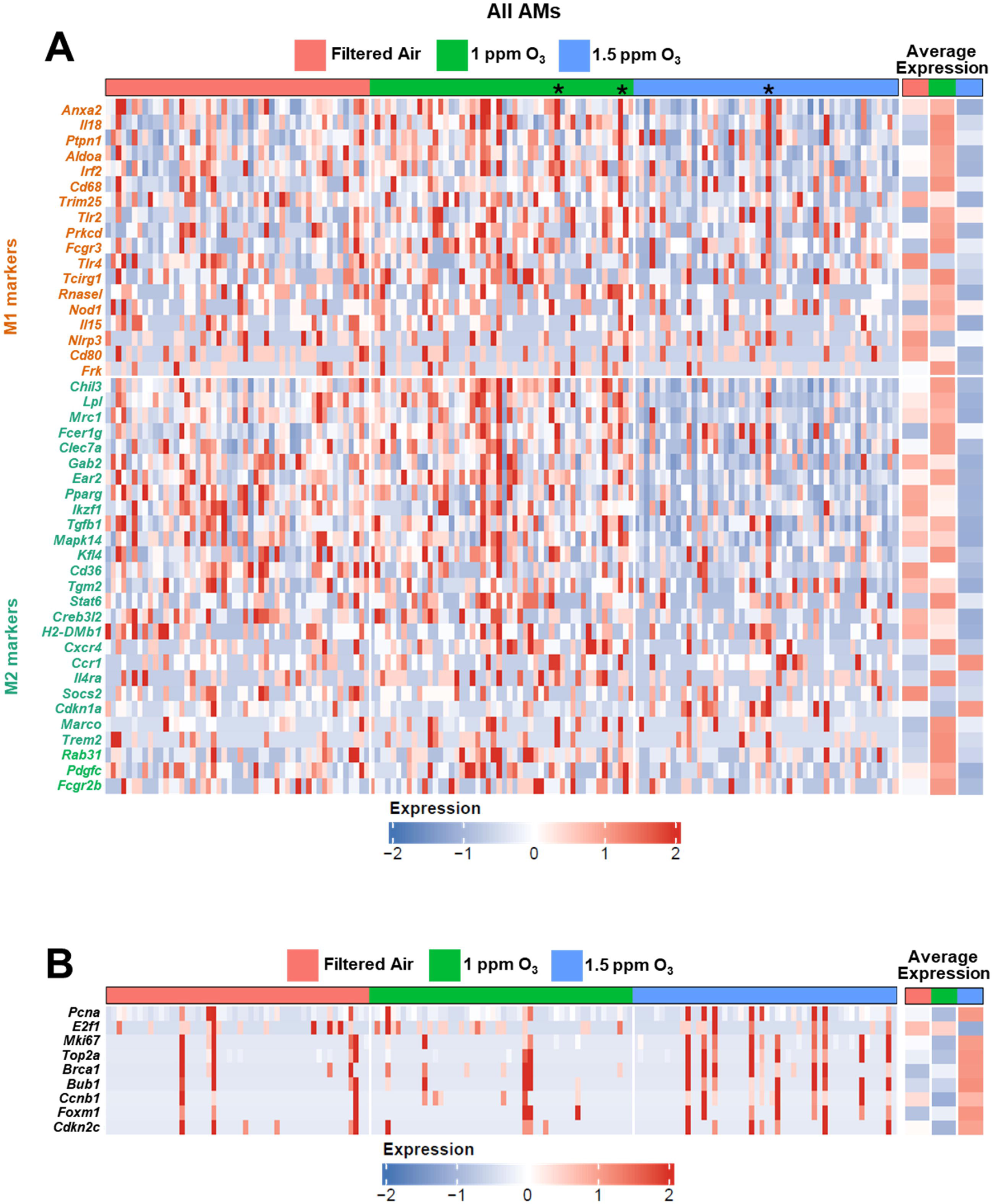
Ozone exposure modulates expression of genes relevant to macrophage activation and cell proliferation. **(A)** Heatmap prepared using normalized gene expression values (z-scores) depicting expression of genes associated with M1 and M2 activation response in 50 randomly-selected AMs from filtered air (FA), 1 ppm ozone (O_3_), and 1.5 ppm O_3_ groups. Asterisks depict cells that display higher copy numbers of transcripts relevant to both M1 and M2 activation responses. (**B**) Heatmap prepared using normalized gene expression values (z-scores) depicting expression of genes associated with proliferative responses in 50 randomly-selected AMs from filtered air (FA), 1 ppm ozone (O_3_), and 1.5 ppm O_3_ groups.

Next, to assess the activation status of AMs from different treatment groups in a cluster-specific manner, we randomly selected 25 cells from each AM cluster (Clusters 0-4) from FA-exposed, 1 ppm O_3_-exposed, and 1.5 ppm O_3_-exposed mice and compared the expression of genes relevant to M1 and M2 activation. In AM cluster 0, compared to FA group, the expression of genes relevant to both M1 and M2 signatures were downregulated in 1 ppm and 1.5 ppm O_3_ groups (**Supplemental Figure S2**). On the other hand, in AM clusters 1 and 2, the M1 and M2 signatures were upregulated in AMs from 1 ppm O_3_ group as compared to FA and 1.5 ppm O_3_ groups **(Supplemental Figure S3, S4**). Next, in AM cluster 3, the M1 and M2 signatures were upregulated in AMs from 1.5 ppm O_3_ group as compared to FA and 1 ppm O_3_ groups (**Supplemental Figure S5**). Lastly, in AM cluster 4, the expression of M1 and M2 signatures did not reveal a definite trend in any treatment groups, indicating that the macrophage activation status in this cluster was minimally affected by O_3_ exposure (**Supplemental Figure S6**).

### Ozone exposure modulates expression of genes associated with proliferative responses

Next, we compared the gene expression of the cell proliferation markers in AMs among FA-exposed, 1 ppm O_3_-exposed, and 1.5 ppm O_3_-exposed mice. Our analyses revealed that the majority of the screened proliferation markers were upregulated in a higher proportion of randomly-selected AMs in 1.5 ppm O_3_ group versus FA and 1 ppm O_3_ groups (**Figure 8B**). Next, we compared the gene expression of the cell proliferation markers in subclusters of AMs among FA-exposed, 1 ppm O_3_-exposed, and 1.5 ppm O_3_-exposed mice. Our analyses revealed that all the proliferation markers, i.e., *Pcna*, *E2f1*, *Mik67*, *Top2a*, *Brca1*, *Bub1*, *Ccnb1*, *Foxm1*, and *Cdkn2c* were expressed in at least 30% of the randomly-selected cluster 3 AMs from at least one of the treatment groups. However, in AM clusters 0,1,2,4, only *Pcna* and *E2f1* genes were expressed in more than 20% of the randomly-selected AMs from at least one of the treatment groups (**Supplemental Figure S7A-C, E**). In AM cluster 0, compared to FA group, the expression of *Pcna* and *E2f1* were downregulated in 1 ppm and 1.5 ppm O_3_ groups (**Supplemental Figure S7A**). In AM cluster 1, the expression of *Pcna* was upregulated in AMs from 1.5 ppm O_3_ group as compared to FA and 1 ppm O_3_ groups, while the expression of *E2f1* was upregulated in 1 ppm O_3_ group as compared to FA and 1.5 ppm O_3_ groups (**Supplemental Figure S7B**). In AM cluster 2, as compared to FA group, the expression of *Pcna* was upregulated in both 1 ppm and 1.5 ppm O_3_ groups, while the expression of *E2f1* was upregulated only in 1 ppm O_3_ group (**Supplemental Figure S7C**). Next, in AM cluster 3, except for *E2f1*, all proliferation markers, i.e., *Pcna*, *Mki67, Top2a, Brca1, Bub1, Ccnb1, Foxm1,* and *Cdkn2c* were upregulated in AMs from 1.5 ppm O_3_-exposed group as compared with FA- and 1 ppm O_3_-exposed groups (**Supplemental figure S7D**). Lastly, in AM cluster 4, the average expression of some proliferation markers, i.e., *Pcna* and *E2f1*, were upregulated in AMs from 1 ppm O_3_-exposed group versus FA-exposed group and some proliferation markers, i.e., *Mki67* and *Cdkn2c*, were upregulated in AMs 1.5 ppm O_3_-exposed group versus FA-exposed group (**Supplemental Figure S7E**).

## Discussion

Inhalation of ambient ozone (O_3_) results in lung inflammation and exacerbates existing respiratory symptoms in susceptible individuals (Liao et al., 2006; Fuller et al., 2022). Previous studies have demonstrated that O_3_ exposure promotes immune cell infiltration, and alters cellular survival and function, including cellular proliferation, activation, and apoptosis (Choudhary, Vo, Paudel, Patial, et al., 2021; Choudhary, Vo, Paudel, Yadav, et al., 2021; Kumagai et al., 2016, 2017). Alveolar macrophages (AMs), a predominant cell type in lung airspaces, represent a highly heterogenous population (Xu-Vanpala et al., 2020; Dick et al., 2022; Wu et al., 2023). Increasing evidence has demonstrated the prevalence of distinct AM subsets in homeostatic (X. Li et al., 2022) and inflammatory conditions (Aran et al., 2019; X. Yu et al., 2024). However, the role of AM heterogeneity in response to O_3_-induced lung inflammation has not been clearly defined. In this study, we employed single-cell RNA sequencing (scRNA-seq) approach to investigate the transcriptomic profile of immune cells, particularly AM, from mice that were exposed to filtered air (FA) or two different concentrations of O_3_. Our findings provided critical insights into the short-term effects of O_3_ exposure on AMs diversity, transcriptomic alterations, and the potential functional modulation.

Immune cell infiltration is a consistent feature of O_3_-induced lung inflammation (Backus et al., 2010; Choudhary, Vo, Paudel, Patial, et al., 2021; Choudhary, Vo, Paudel, Yadav, et al., 2021; Gabehart et al., 2014; Guttenberg et al., 2024; Mathews et al., 2018; Oakes et al., 2013; Tovar et al., 2020; Verhein et al., 2015). Consistent with these previous reports, we observed the infiltration of neutrophils in the lung airspaces of O_3_-exposed mice as compared with FA-exposed mice. Consistent with the increase recruitment of neutrophils in O_3_-exposed mice, the BALF levels of neutrophil-specific chemokines, i.e., MIP-2, KC, and LIX were found elevated. The annotated cellular clustering revealed an increase in other immune cells, e.g., CD4+ Tcells, CD4+/CD8+ Tcells, and type 2 innate lymphoid cells (ILC2s) in 1.5 ppm O_3_-exposed mice as compared with 1 ppm O_3_-exposed mice, highlighting the concentration-dependent response to O_3_ exposure in altering immune cell composition in the lung airspaces. The increase in Th2 inflammation-relevant immune cells in 1.5 ppm O_3_ as compared with 1 ppm O_3_ was consistent with the significant increase in MDC/CCL22 levels in the bronchoalveolar lavage fluid (BALF) of 1.5 ppm O_3_-exposed mice versus 1 ppm O_3_-exposed mice. Despite the enhanced recruitment of other immune cells in the O_3_-exposed mice as compared with the FA-exposed mice, AMs remained the predominant cell type in the airspaces of mice in all treatment groups. Therefore, we compared the gene expression profiles of AMs between 1 ppm O_3_-exposed mice versus FA and 1.5 ppm O_3_-exposed mice versus FA.

The expansion of macrophage population through local proliferation and/or recruitment is a common response observed in airspace stress caused by various intrinsic factors as well as extrinsic agents (Jenkins et al., 2011; Z. Liu et al., 2019; van de Laar et al., 2016; Morse et al., 2019; Aegerter et al., 2020; Machiels et al., 2017; Misharin et al., 2017). While the recruitment of inflammatory macrophages, expressing gene signatures distinct from resident AMs, takes place in sustained and prolonged infection or exposure to a stress (Aegerter et al., 2020; Machiels et al., 2017; Misharin et al., 2017), the local expansion of resident AMs occurs as a short-term response to a lower degree of stress (Hotchkiss et al., 1989; Jenkins et al., 2011; Z. Liu et al., 2019). Our analyses revealed concentration-dependent and AM cluster-specific proliferative responses. Our findings are consistent with a previous study on O_3_-exposed rats, where proliferative macrophages were the primary cause of increase in AM counts in earlier stages post-exposure (Hotchkiss et al., 1989; Tredaniel et al., 1994). The increased AM proliferative signatures in 1.5 ppm group were also associated with significantly higher M-CSF levels in BALF. Of note, M-CSF is a potent proliferation inducer, which is eleased by various lung cells, including macrophages, neutrophils, fibroblasts and endothelial cells (Braza et al., 2018; Clinton et al., 1992; Falkenburg et al., 1990; Tang et al., 2018). The exact source of M-CSF and other proliferation triggers and their role in the expansion of AM population remain unclear.

Previous studies have demonstrated that AMs remain functionally restrained at homeostasis (Hussell & Bell, 2014), but become functionally activated upon exposure to O_3_ (Backus et al., 2010; Choudhary, Vo, Paudel, Patial, et al., 2021; Choudhary, Vo, Paudel, Wen, et al., 2021; Fakhrzadeh et al., 2004; Pendino et al., 1994; Reinhart et al., 1999; Sunil et al., 2012). Consistent with these reports, we observed a markedly increased number of differentially-expressed genes (DEGs) and corresponding enriched biological pathways between O_3_-exposed mice versus FA-exposed mice, suggesting the effect of O_3_ exposure in modulating gene expression and biological functions of AMs. Additionally, the alteration in the transcriptome and biological functions of AMs from 1.5 ppm O_3_-exposed mice versus 1 ppm O_3_-exposed mice further emphasized the concentration-dependent effects of O_3_ exposure. Acute exposure to O_3_ resulted in enhanced production of interleukin-1, apoptosis, and activation of transcription factor NF-kB (Pendino et al., 1994; Sunil et al., 2012). Consistent with the previous reports, the ingenuity pathway analyses (IPA) results highlighted the common pathways in both 1 ppm and 1.5 ppm O_3_-exposed AMs, including interleukin-1 family signaling, oxidative phosphorylation, non-canonical NF-kB signaling, and regulation of apoptosis. In addition, we also observed the activation of common pathways between 1 ppm and 1.5 ppm O_3_-exposed AMs, including cellular response to hypoxia and KEAP1-NFE2L2 pathways, which were only reported to be elevated in chronic O_3_ exposure (Wiegman et al., 2014).

Macrophage activation, i.e., enhanced functioning, is a typical response that has been widely investigated (Groves et al., 2013; Mathews et al., 2015). On the other hand, several acute studies using high O_3_ concentration exposure (1.5 ppm – 2 ppm) demonstrated poor phagocytic abilities of AMs (Jakab & Hemenway, 1994; Mikerov, Gan, et al., 2008; Mikerov, Haque, et al., 2008), suggesting that high O_3_ concentration compromises AM functions. Moreover, AMs have been identified as targets of O_3_-induced oxidative DNA damage (Sunil et al., 2012), which has been associated with impaired macrophage functions, including defects in proliferation, delayed differentiation, and increased senescence. and an enhanced airway inflammatory response (Luo et al., 2018; Pereira-Lopes et al., 2015). Consistent with these studies, our IPA data showed that AMs from the 1.5 ppm O_3_-exposed mice were uniquely activated in multiple DNA repair, DNA damage bypass, and cell cycle checkpoints pathways. Moreover, it has been reported previously that macrophages exhibit dysregulated inflammatory response and altered immune functions in the absence of proper lipid metabolic programming (Lee & Bensinger, 2022). In this current study, the regulation of lipid metabolism pathway was suppressed in AMs from cluster 3 of the 1.5 ppm O_3_-exposed mice. We speculated that 1.5 ppm O_3_ concentration disrupted the tightly controlled cell cycle checkpoints by bypassing the DNA damage, thus promoting irregular cell proliferation. We further speculated that the newly proliferated AMs were functionally less mature. However, further experimentation is needed to test these hypotheses.

Several reports indicate that macrophages can reprogram their cholesterol biosynthetic machinery to aid in host defense responses (Blanc et al., 2013; Hayakawa et al., 2022; Lee & Bensinger, 2022; Madenspacher et al., 2020). Previous reports have shown upregulation of genes relevant to the cholesterol biosynthesis pathways in the lung following repetitive O_3_ exposure (Cho et al., 2021; Choudhary, Vo, Paudel, Patial, et al., 2021; Colonna, 2015; Tovar et al., 2020). Consistent with previous studies, in the current study, despite one-time exposure paradigm, we found enrichment of pathways driven by cholesterol biosynthesis relevant genes (*Cyp51a1, Dhcr24, Fdft1, Fdps, Hmgcr, Hmgcs1, Idi1, Msmo1, Pmvk, Sc5d, Sqle*). Oxysterols, e.g., SecoA, SecoB, and [3-epoxycholesterols, are known to be upregulated in O_3_-exposed airways, and are implicated as potential mediators of O_3_ toxicity (Pryor et al., 1992; Pulfer et al., 2005; Pulfer & Murphy, 2004; Speen et al., 2016). Another report by Duffney et al demonstrated that oxysterols can form adducts with CD64 and CD206 which can interfere with the phagocytic potential of AMs (Duffney et al., 2020). In contrast, cholesterol derivatives are also found to be beneficial in host responses. For instance, Madenspacher et al studied the role of 25 hydroxycholesterol (25HC), a product of an important cholesterol biosynthesis enzyme (cholesterol-25-hydroxylase), in LPS-induced inflammation (Madenspacher et al., 2020). They demonstrated that 25HC prevent lipid overload in AMs, in an anti-inflammatory nuclear receptor liver-x-receptor (LXR)-dependent manner, which is critical for the Mertk-mediated efferocytotic removal of neutrophils and resolution of inflammation in LPS-challenged mice (Madenspacher et al., 2020). It is likely that the cholesterol derivatives are playing similar role in the efferocytosis of O_3_-induced neutrophils and subsequent resolution of inflammation. It is also likely that the upregulated cholesterol biosynthesis pathways in macrophages serve as a source of cholesterol derivatives in the O_3_-exposed airspaces. However, the exact identity and source of the O_3_-induced triggers for the reprogramming of cholesterol biosynthesis in AMs, and their functional impact on immune and epithelial cells remain unknown and require further investigation.

Mitochondria are the major source of reactive oxygen species (ROS) and free radicals during oxidative phosphorylation (Boveris et al., 1972; Boveris & Chance, 1973), and play a key role in lipid metabolism and cholesterol homeostasis (Han et al., 2021; L. Li et al., 2022; Torres et al., 2021). Variety of environmental insults result in overproduction of mitochondrial ROS (mtROS), which could cause mitochondrial dysfunction, i.e., damage mitochondrial protein, mitochondrial lipid membrane, and mitochondrial DNA (Brookes et al., 2004). Increasing evidence supports the important role of mitochondrial dysfunction in the development and pathogenesis of lung diseases (Aguilera-Aguirre et al., 2009; Jablonski et al., 2017; Sharma et al., 2021; J. Yu et al., 2016). Moreover, the O_3_-induced mitochondrial structural damage was previously reported in epithelial cells (Lopez et al., 2007), cardiomyocytes (Tian et al., 2021), airway smooth muscle cells (Wiegman et al., 2015), and microglia (Valdez et al., 2020). In the current study, pathways related to mitochondrial responses, e.g., oxidative phosphorylation, respiratory electron transport, mitochondrial protein import, mitochondrial dysfunction, and complex I biogenesis, were enriched in the AMs from 1.5 ppm O_3_-exposed group versus the 1 ppm O_3_-exposed and FA groups. However, the exact mechanism and consequences of O_3_-induced mitochondrial dysfunction in AMs remains unclear and requires further experimentation.

Translational control is critical for cellular preservation in response to stress (Holcik & Sonenberg, 2005). One of the key mechanisms of translational control during stress is the phosphorylation of eIF2, which results in the transcriptional activation of stress response genes (Harding et al., 2000). In the current study, we found enrichment of protein translation pathways driven by eIF2 signaling and other relevant genes, e.g., *Eif3f, Eif3h, Rpl21, Rpl23, Rpl26, Rpl27a, Rpl28, Rpl34, Rpl36, Rpl36a, Rpl37, Rpl4, Rps11, Rps15a, Rps19, Rps26, Rps27,* and *Rps8*. Additionally, previous studies suggested the involvement of eIF2 signaling in antibacterial response (S. Liu et al., 2007; Nakayama et al., 2010; Shrestha et al., 2012; Woo et al., 2009). For instance, using three very diverse pathogens, e.g., *Yersinia pseudotuberculosis, Listeria monocytogenes*, and *Chlamydia trachomatis*, Shrestha et al has demonstrated the direct linkage between eIF2 signaling and the antibacterial defense system (Shrestha et al., 2012), suggesting the roles of eIF2 signaling pathway in the host defense mechanism.

Free fatty acid receptor 2 (*Ffar2*/*Gpr43*) is widely expressed in various inflammatory cells such as macrophages and neutrophils (Kamp et al., 2016; McNelis et al., 2015). The reported roles of *Ffar2* in inflammation were controversial. For instance, *Ffar2* gene deletion was reported to increase inflammation in arthritis model and allergic airway model (Maslowski et al., 2009). On the other hand, *Ffar2* gene deletion was reported to decrease inflammation in ethanol-induced inflammation model (M. H. Kim et al., 2013), and dextran sulfate sodium (DSS)-induced colitis model (Sina et al., 2009). Further, deletion of *Ffar2* was reported to increase perivascular fibrosis and increase cancer progression. In the current study, *Ffar2* was commonly increased in both 1 ppm and 1.5 ppm O_3_-exposed groups, suggesting the inflammatory role of *Ffar2* in O_3_-induced lung injury. However, further experimentation is warranted to elucidate whether the elevated *Ffar2* expression was associated with pro-inflammatory or anti-inflammatory responses.

Gene signatures associated with macrophages could indicate the nature of their immediate microenvironment. Macrophages may respond to products released from epithelial cells upon injury by O_3_ resulting in classical macrophage activation (M1) and release products such as reactive oxygen species (ROS) and TNF-α, which further promote lung injury (Cho et al., 2001; Fakhrzadeh et al., 2004; Pendino et al., 1995). On the other hand, another study has shown the presence of alternatively activated AMs (M2) in the O_3_-exposed lungs (Mathews et al., 2015). The M2 macrophages release anti-inflammatory mediators such asinterleukin-10 (IL-10) and actively participate in wound repair, clearance of apoptotic/necrotic cells (Dahl et al., 2007; Ishii et al., 1998), and resolution of inflammation (Backus et al., 2010; Reinhart et al., 1999). We observed interesting expression readouts of macrophage activation-associated gene signatures. The assessment of macrophage-activation relevant gene signatures in the entire AM population revealed that exposure to 1 ppm O_3_ causes an upregulated expression of genes associated with M1 as well as M2 activation. In contrast, as compared to both FA- and 1ppm O_3 -_exposure groups, AMs from the 1.5 ppm O_3_-exposed mice exhibited downregulated expression of genes relevant to both M1 and M2 activation. We found interesting patterns of gene signatures associated with alternative and classical AMs activation responses in a cluster-specific manner. Particularly, analyses of gene expression for AM in cluster 3 suggest that the 1.5 ppm O_3_ exposure results in upregulated expression of genes associated with M1 as well as M2 activation. On the other hand, analyses of gene expression for AM from clusters 1 and 2 suggest that 1.5 ppm O_3_ exposure downregulates the expression of genes associated with M1 as well as M2 activation. A previous report indicated the co-existence of both M1 and M2 gene signatures in AMs retrieved from BALF of C57BL/6J mice that were sub-chronically exposed to 0.8 ppm O_3_ (Choudhary, Vo, Paudel, Patial, et al., 2021). However, it remained unclear whether the mixed M1/M2 signature was the result of population heterogeneity or cellular plasticity. In this study, we assessed the M1 and M2 signatures in all AM clusters from FA-exposed, 1 ppm O_3_-exposed and 1.5 ppm O_3_-exposed mice. Interestingly, we found elevated levels of both M1 and M2 markers within the same cell suggesting that the expression of gene signatures associated with M1 and M2 activations is not mutually exclusive.

This study revealed some interesting findings. First, O_3_ exposure results in a concentration-dependent increase in the levels of inflammatory mediators in the airspaces of mice. Second, within 24 h of O_3_ exposure, the AM population diversity remains unchanged, suggesting minimal recruitment of inflammatory macrophage populations. Third, DEG analyses revealed greater perturbation in gene expression in AMs from the 1.5 ppm versus the 1 ppm O_3_-exposed group. Fourth, exposure to higher O_3_ concentration is associated with the enrichment of pathways involved in cell cycle checkpoint, DNA damage and repair, cholesterol biosynthesis, and mitochondrial responses. Finally, assessment of macrophage activation status revealed the concentration-dependent expression of genes relevant to macrophage activation in a cluster-specific manner. Taken together, our data highlight the intrinsic transcriptomic disturbances in highly heterogenous alveolar macrophage subpopulations in murine airspaces post O_3_ exposure at single-cell resolution level.

## Acknowledgment

We thank Amber McCoy and Gabrielle Cannon from the UNC Advanced Analytic Core (UNC Center for GI Biology and Disease, P30 DK034987) for their assistance with the 10x Genomics scRNA-Seq library preparation and analysis. We thank Michael Vernon at the UNC High Throughput Sequencing Facility for the technical support. This facility is supported by the University Cancer Research Fund, Comprehensive Cancer Center Core Support grant (P30-CA016086), and UNC Center for Mental Health and Susceptibility grant (P30-ES010126).

## Supplemental Figure Legends

**Supplemental Figure S1: (A)** Integrated UMAP combined from filtered air (FA) (solid pink dots), 1 ppm ozone (O_3_) (solid green dots) and 1.5 ppm O_3_ (solid blue dots) groups**. (B)** Integrated UMAP split between FA (solid pink dots), 1 ppm ozone (O_3_) (solid green dots) and 1.5 ppm O_3_ (solid blue dots) groups.

**Supplemental Figure S2:** Heatmap prepared using normalized gene expression values (z-scores) depicting expression of genes associated with M1 and M2 activation in 25 randomly-selected AMs in cluster 0 from filtered air (FA), 1 ppm ozone (O_3_), and 1.5 ppm O_3_ groups.

**Supplemental Figure S3:** Heatmap prepared using normalized gene expression values (z-scores) depicting expression of genes associated with M1 and M2 activation in 25 randomly-selected AMs in cluster 1 from filtered air (FA), 1 ppm ozone (O_3_), and 1.5 ppm O_3_ groups.

**Supplemental Figure S4:** Heatmap prepared using normalized gene expression values (z-scores) depicting expression of genes associated with M1 and M2 activation in 25 randomly-selected AMs in cluster 2 from filtered air (FA), 1 ppm ozone (O_3_), and 1.5 ppm O_3_ groups.

**Supplemental Figure S5:** Heatmap prepared using normalized gene expression values (z-scores) depicting expression of genes associated with M1 and M2 activation in 25 randomly-selected AMs from cluster 3 in filtered air (FA), 1 ppm ozone (O_3_), and 1.5 ppm O_3_ groups.

**Supplemental Figure S6:** Heatmap prepared using normalized gene expression values (z-scores) depicting expression of genes associated with M1 and M2 activation in 25 randomly-selected AMs in cluster 4 from filtered air (FA), 1 ppm ozone (O_3_), and 1.5 ppm O_3_ groups.

**Supplemental Figure S7:** Heatmap prepared using normalized gene expression values (z-scores) depicting expression of genes associated with proliferative responses in 25 randomly-selected AMs from filtered air (FA), 1 ppm ozone (O_3_), and 1.5 ppm O_3_ groups in (**A**) AM cluster 0, (**B**) AM cluster 1, (**C**) AM cluster 2, (**D**) AM cluster 3, and (**E**) AM cluster 4.

