## Supplementary figures and images for "Single-cell transcriptomics reveal alveolar macrophages-specific responses in single-hit ozone exposure model in mice"

### Supplemental Figure 1

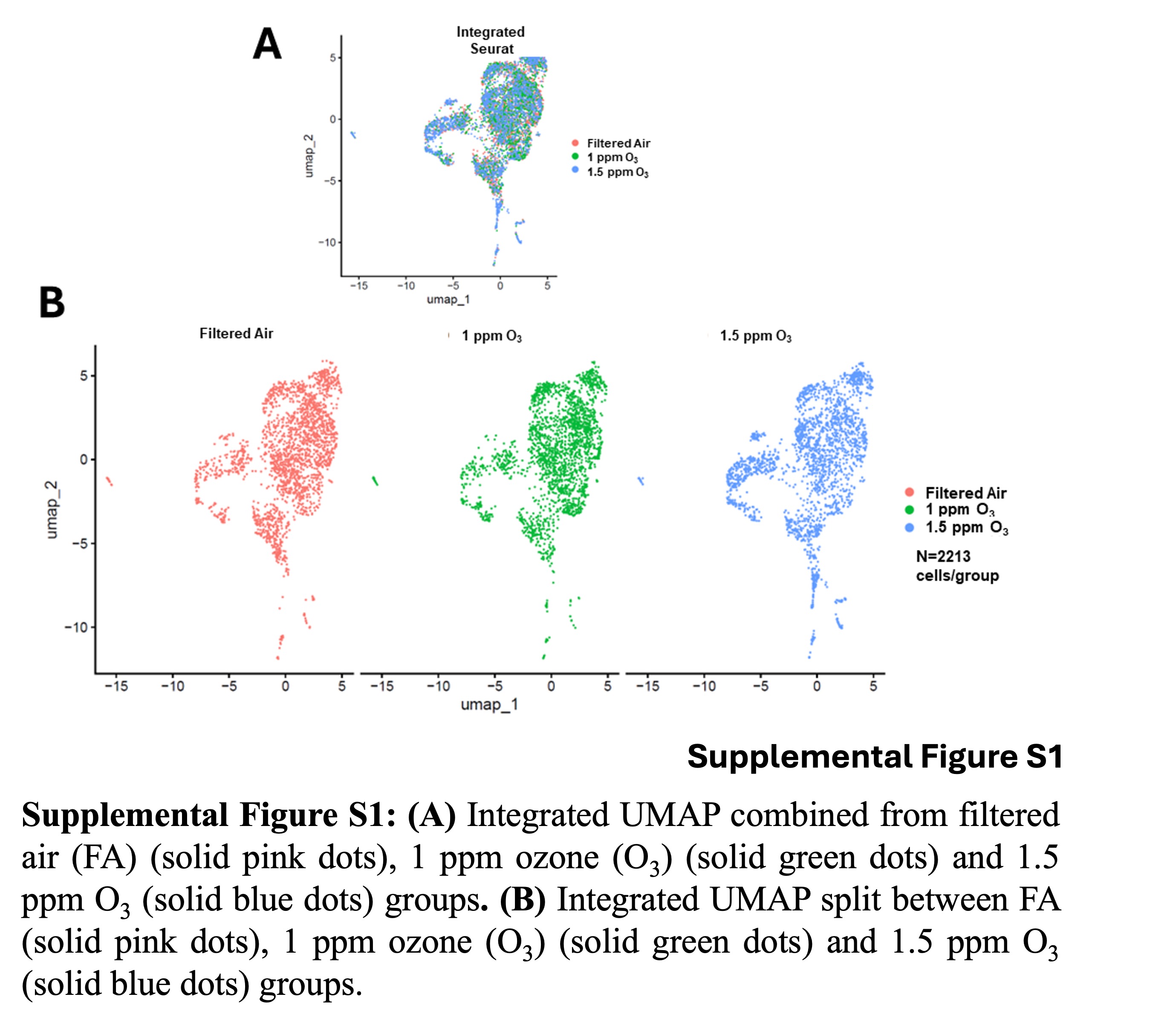

### Supplemental Figure 2

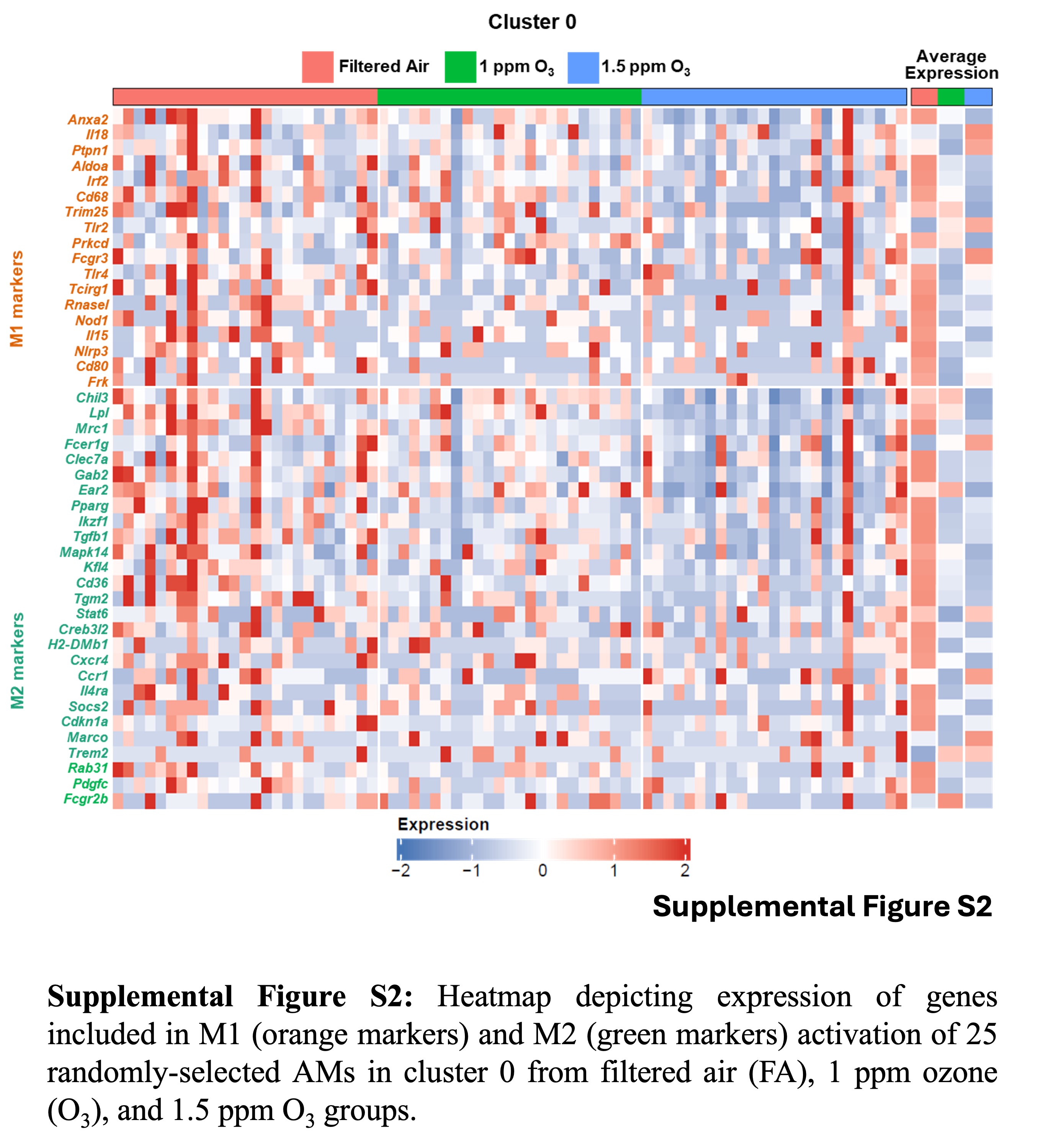

### Supplemental Figure 3

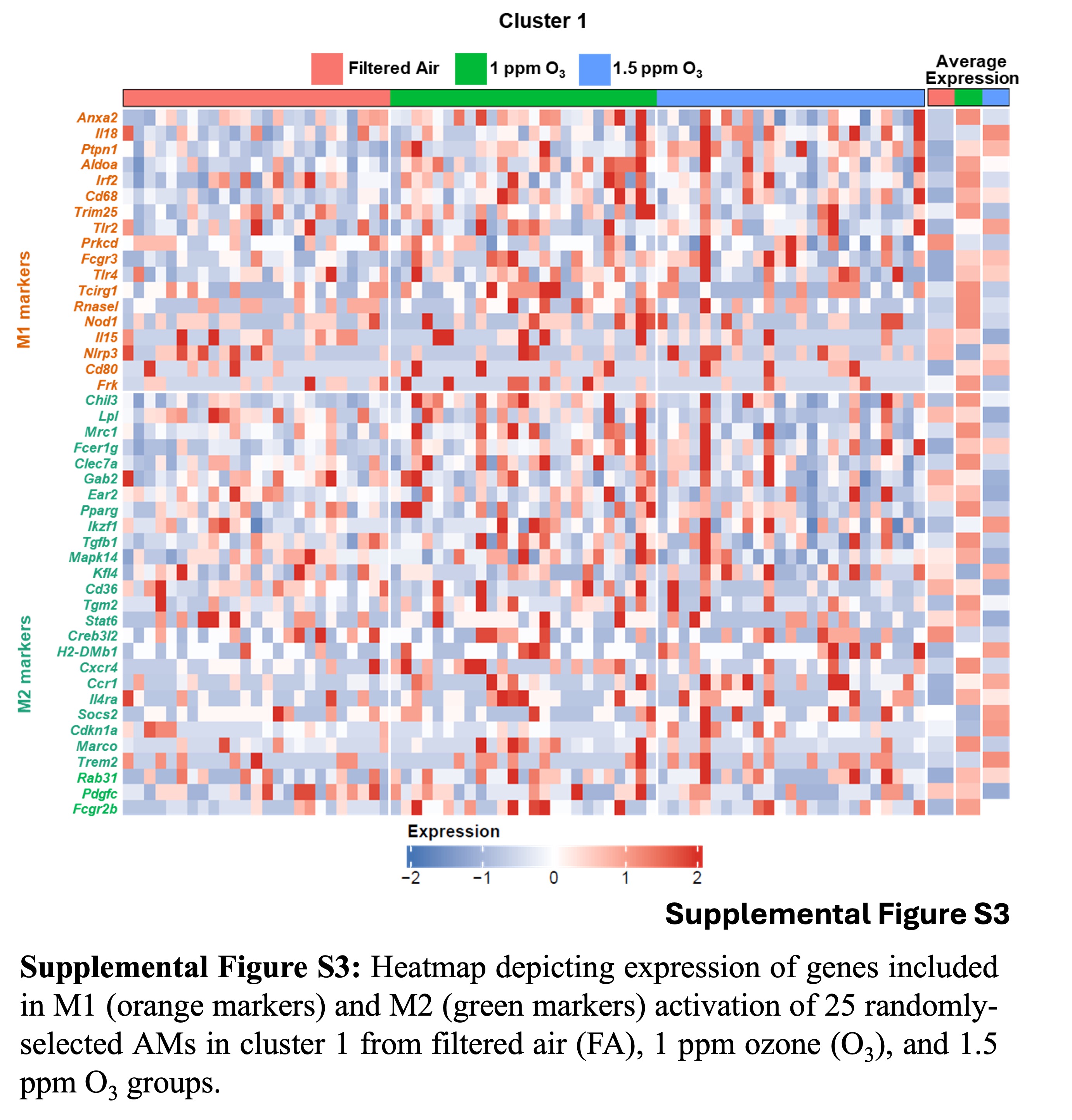

### Supplemental Figure 4

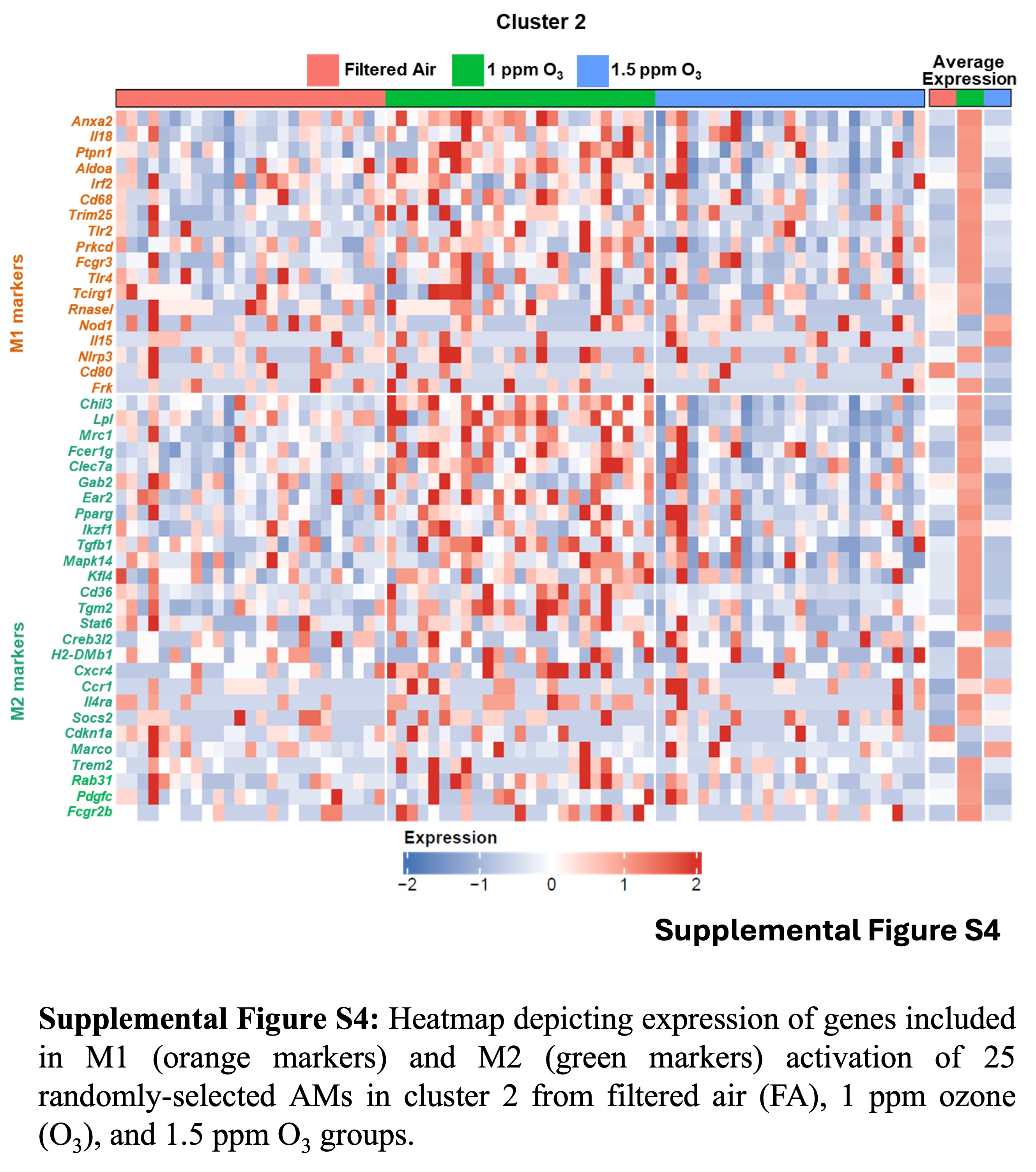

### Supplemental Figure 5

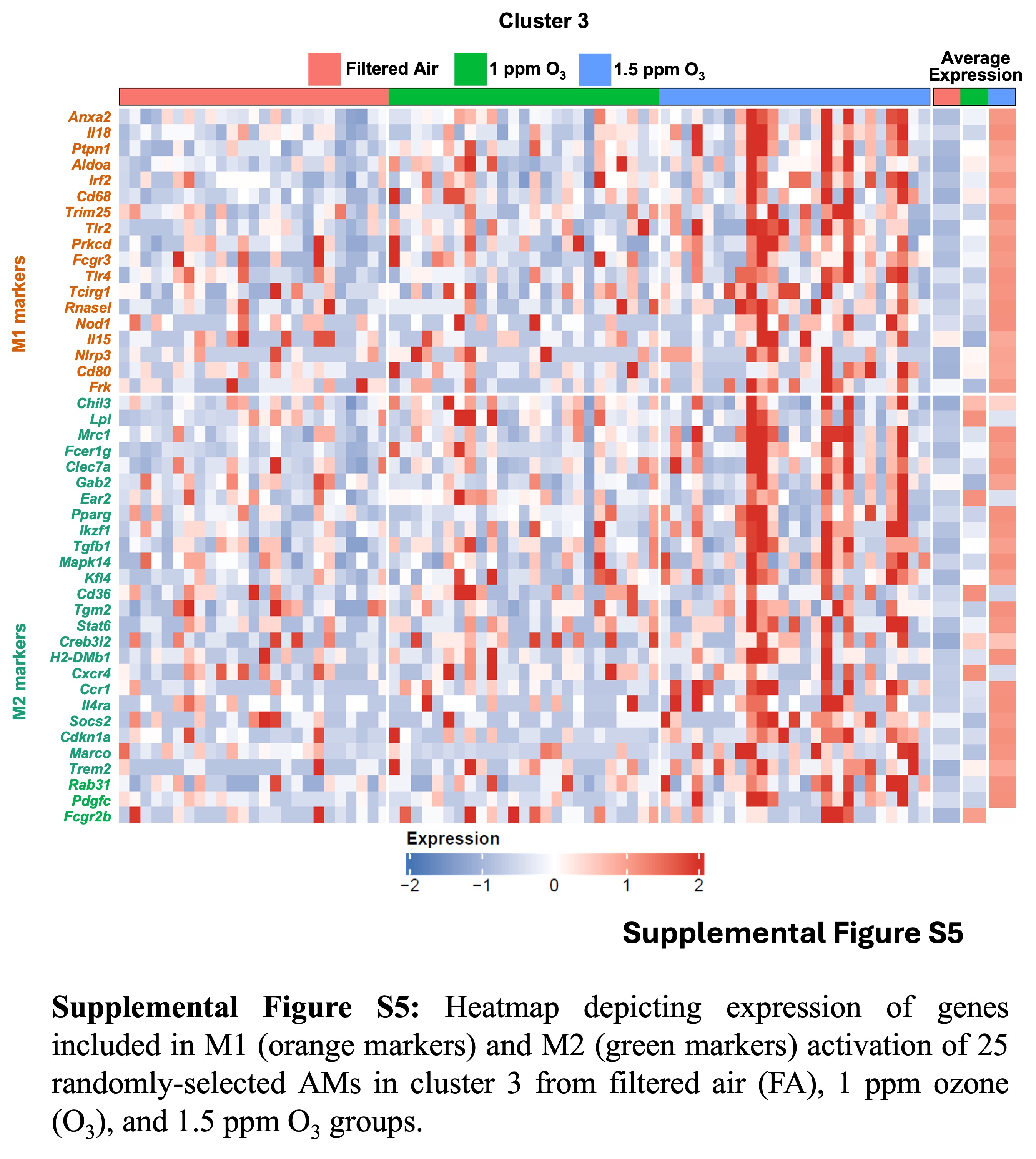

### Supplemental Figure 6

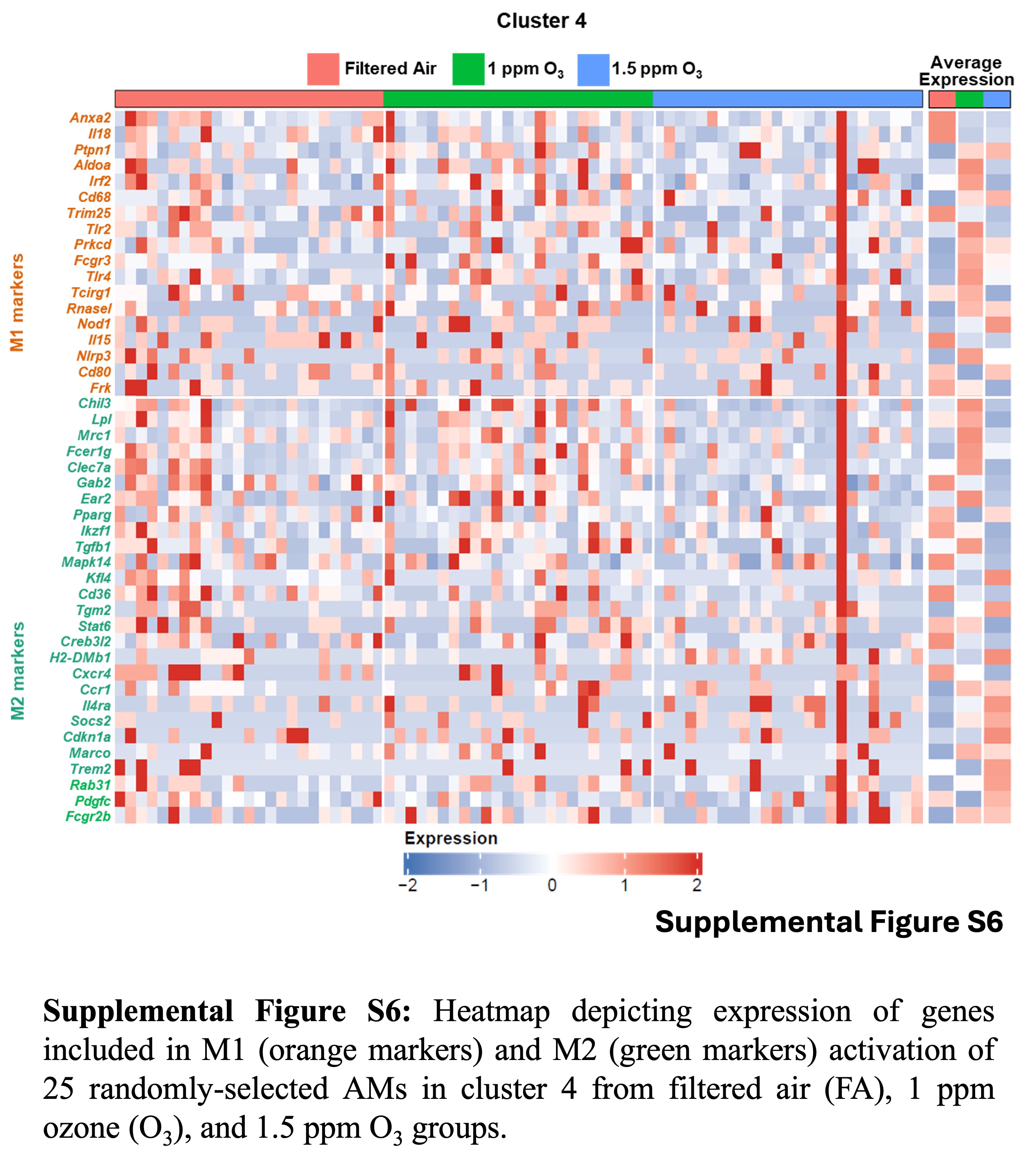

### Supplemental Figure 7

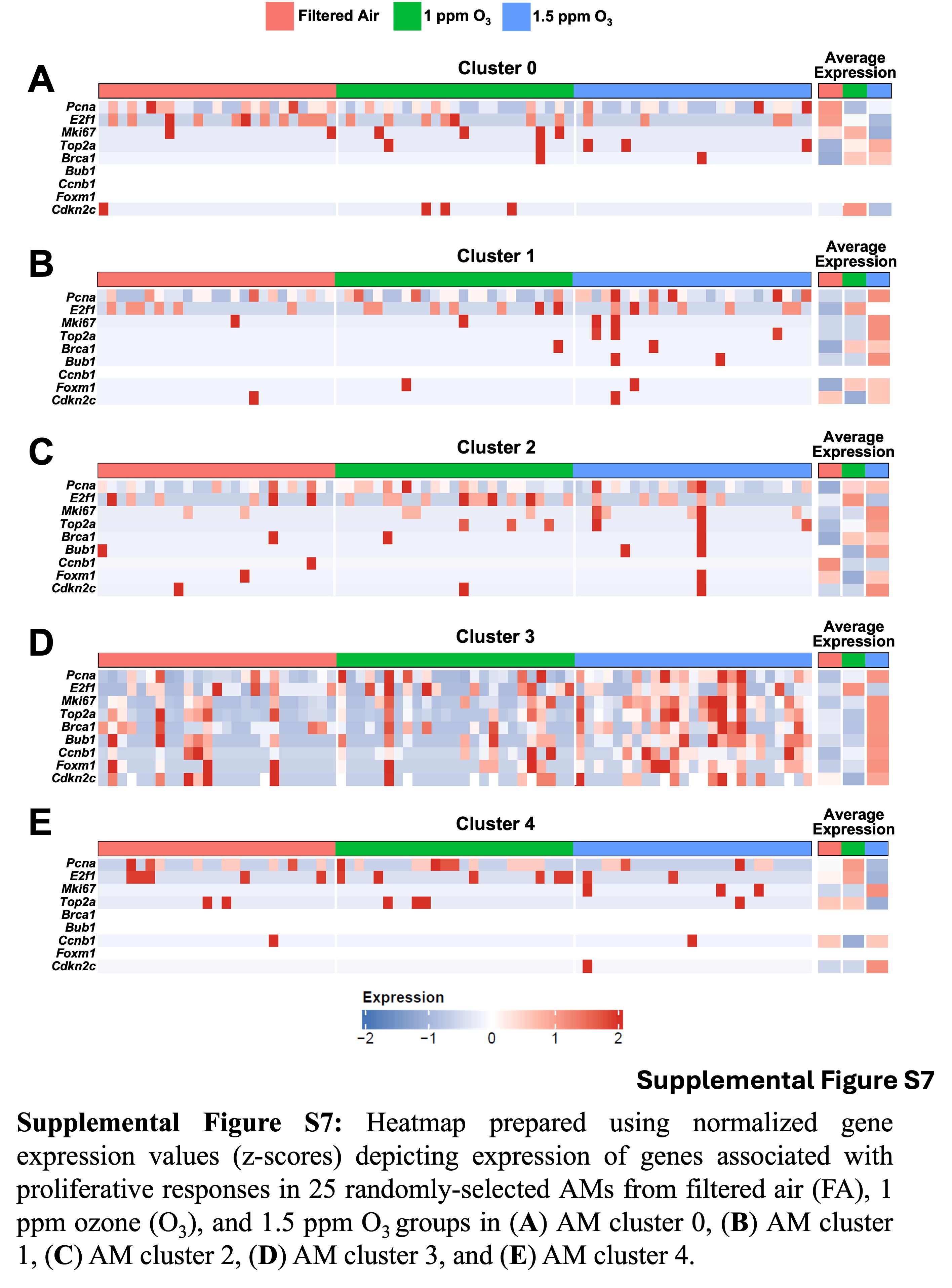
